# Vancomycin resistance in *Enterococcus faecium* from the Dallas, Texas area is conferred predominantly on pRUM-like plasmids

**DOI:** 10.1101/2021.02.16.431552

**Authors:** Moutusee Islam, Belle Sharon, Ada Abaragu, Harita Sistu, Ronda L. Akins, Kelli Palmer

**Affiliations:** Department of Biological Sciences, University of Texas at Dallas, Richardson, Texas, USA; Methodist Health System, Dallas, Texas, USA

**Keywords:** *Enterococcus*, mobile genetic element, plasmid, vancomycin resistance

## Abstract

Vancomycin-resistant *E. faecium* (VREfm) is a significant public health concern because of limited treatment options. Genomic surveillance can be used to monitor VREfm transmission and evolution. Genomic analysis of VREfm has not been reported for the Dallas/Fort Worth/Arlington, Texas, area, which is currently the 4th largest metropolitan area in the United States. Our study aimed to address this gap in knowledge by analyzing the genomes of 46 VREfm and one vancomycin-sensitive comparator collected during routine fecal surveillance of high-risk patients upon admission to a Dallas, Texas hospital system (August to October, 2015). 31 complete and 16 draft genome sequences were generated. The closed VREfm genomes possessed up to 12 extrachromosomal elements each. Overall, 251 closed putative plasmid sequences assigned to previously described and newly defined *rep* family types were obtained. Phylogenetic analysis identified 10 different sequence types (STs) among the isolates, with the most prevalent being ST17 and ST18. Strikingly, all but three of the VREfm isolates encoded *vanA*-type vancomycin resistance within Tn*1546-* like elements on a pRUM-like (*rep17*) plasmid backbone. Relative to a previously reported typing scheme for the *vanA*-encoding Tn*1546*, new variants of the Tn*1546* were identified that harbored a combination of 7 insertion sequences (IS), including 3 novel IS elements reported in this study (*ISEfa16, ISEfa17* and *ISEfa18*). We conclude that pRUM-like plasmids are important vectors for vancomycin resistance in the Dallas, Texas area and should be a focus of plasmid surveillance efforts.

**Importance:** Vancomycin is an antibiotic used to treat infections caused by multidrug-resistant Gram-positive bacteria. Vancomycin resistance is common in clinical isolates of the Gram-positive pathogen *Enterococcus faecium*. In *E. faecium*, vancomycin resistance genes can be disseminated by plasmids with different host ranges and transfer efficiencies. Surveillance of resistance plasmids is critical to understanding antibiotic resistance transmission. This study analyzed the genome sequences of VREfm collected from the Dallas, Texas area, with particular focus on the mobile elements associated with vancomycin resistance genes. We find that a single plasmid family, the pRUM-like family, is associated with vancomycin resistance in the majority of isolates sampled. Our work suggests that the pRUM-like plasmids should continue to be studied to understand their mechanisms of maintenance, transmission, and evolution in VREfm.

## Introduction

Originally described as a commensals of the healthy human gut, the enterococci attracted attention in the 1990s due to their alarming resistance to multiple antibiotics (1). Vancomycin-resistant enterococci (VRE) were first reported in 1988 in Europe (2) and in the USA in 1989 (3) and are a major concern (4). Vancomycin-resistant *E. faecium* (VREfm) is particularly concerning due to limited treatment options and widespread occurrence of vancomycin resistance among infection isolates of *E. faecium* (4).

Phylogenetic analysis of *E. faecium* strains revealed the presence of two major clades, referred to as Clade A and B, with some evidence for subclades A1 and A2 within Clade A (5, 6). Clade B *E. faecium* strains are mostly commensals of the healthy human gut. The multidrug-resistant *E. faecium* strains responsible for hospital outbreaks typically belong to Clade A1, whereas animal gut commensals typically belong to Clade A2 (6). Multi-locus sequence typing (MLST) pre-dates whole genome-based phylogenetic analyses and has been used globally to classify *E. faecium* isolates by nucleotide sequence variations occurring in 7 housekeeping genes (7). Isolates are assigned to different sequence types (ST) based on the allelic variants. Most reported hospital outbreaks of VREfm have emerged from a single genetic lineage, referred to as clonal complex-17 (CC17), that was founded by the sequence type ST17 (8).

Vancomycin resistance genes are encoded within mobile elements in VREfm and can be horizontally disseminated (9–11). VanA-type resistance commonly occurs among VREfm, and VanA-type resistance genes are typically encoded within Tn*1546* (12). Tn*1546* may be chromosomally integrated or plasmid-borne. Different plasmid backbones carrying *Tn1546*, including Inc18, pRUM, and pLG1, have been reported worldwide (for *e.g*. references (10, 13)). Structural variations of Tn*1546* due to the presence of insertion sequences (IS) including *IS1251, IS1216, IS1485* and *ISEf1* have also been reported. These previous studies characterized Tn*1546* structural variation by PCR mapping and then sequencing of the overlapping fragments (10, 13, 14). However, in the absence of a completely closed genome assembly, conclusive links between *vanA, Tn1546* variants, and specific plasmid backbones in VREfm can be difficult to achieve.

Surveillance of antibiotic resistance elements is important because changes in their host range and transfer frequency could evolve, which could impact clinical care. Utilization of whole genome sequencing for epidemiologic study provides an opportunity for multilevel analysis that includes detection of novel mobile genetic elements and monitoring DNA sequence changes in resistance gene vectors. The advent of long-read next generation sequencing (NGS) by Pacific Biosciences and Oxford Nanopore Technology (ONT) has made it possible to generate genome assemblies of a large number of bacterial isolates in a cost-effective manner. The combination of long (ONT in this study) and short (typically Illumina) reads generates high quality, completely closed genome assemblies with fully assembled mobile genetic elements (15).

In this study, we analyzed phylogenetic relationships and plasmid diversity among a previously reported collection of VREfm from the Dallas, Texas area (16). These isolates were recovered from rectal surveillance of high-risk patients upon admission to a Dallas hospital system. These represent colonization reservoirs from which future hospital outbreaks could emerge. Some results of our study are expected, including the predominance of CC17/Clade A1 among the VREfm isolates. Other results highlight important areas for future study. Specifically, we note the high number of extrachromosomal elements in VREfm, with some isolates possessing 12 unique elements independent from the chromosome. How these multiple elements are coordinated and maintained in VREfm, and what they contribute to VREfm physiology, are largely unknown and are important topics for future investigation. Moreover, pRUM-like plasmids are the dominant carriers of vancomycin resistance genes among these VREfm. This plasmid family should continue to be investigated in laboratory experiments to understand its maintenance and transmission mechanisms.

## Materials and Methods

### Bacterial strains analyzed in this study

The isolates used in this study (**Dataset S1A**) were cultured from surveillance rectal swabs of high-risk patients on hospital admission (August 2015 to October 2015). Their clinical collection and initial characterization was previously reported (16) and is briefly summarized here. Rectal swabs from patients admitted with at least one of the following risk factors (hospitalization for ≥2 consecutive nights in the preceding 30 days, transferred from another medical facility, residence in a nursing home or extended/long term care facility, or the presence of decubitus ulcer or a draining wound) were cultured on Spectra VRE (Remel) agar as a part of routine infection control surveillance. Instead of disposal after clinical procedures were complete, cultures positive for presumptive VRE growth were coded numerically, de-identified, and transferred to the University of Texas at Dallas for further analysis. Isolates were coded by de-identified patient number. If multiple colonies were collected from a Spectra VRE plate, this was indicated by hyphenated numerics. For example, the VRE isolates 124-1, 124-2 and 124-3 were collected from the same rectal swab sample from the same patient (de-identified patient #124). No patients were sampled longitudinally.

In our previous study of these isolates, presumptive VREfm were confirmed by *ddl* typing and for the presence of *vanA* or *vanB* genes by PCR (16, 17). All isolates used in this study, with the exception of 163-1, initially grew on brain heart infusion (BHI) agar supplemented with 256 μg/mL vancomycin. The vancomycin MIC of isolate 163-1 was previously determined to be 2 μg/mL by broth microdilution (16). The vancomycin MIC of 10 additional isolates were previously determined to be >256 μg/mL by broth microdilution (16). The vancomycin MIC of all other isolates were determined in this study using the same broth microdilution method previously reported (16).

### Genome sequencing and assembly

Methods for DNA isolation, MinION sequencing, Illumina sequencing, and hybrid genome assembly are described in the Supplemental Text.

### Phylogenetic analyses and MLST

Genome assemblies were annotated with PROKKA (v 1.12) (18). A sequence alignment was generated for 1706 core genes by ROARY (v 3.12) (19) using PRANK where the paralogs were not split. A phylogenetic tree was built by RAxML (v 8.2.12) (20) with rapid bootstrapping (1000 inferences) using the GTR+GAMMA model and subsequent ML search on the core gene alignment where the outgroup was specified to be the reference genome *E. faecium* 1,231,501 (GenBank accession number ACAY01). The phylogenetic tree was visualized with iTOL (v4) (21). MLST was determined by the *E. faecium* MLST database (https://pubmlst.org/efaecium/) (7, 22). The novel sequence type ST1703 and novel *gyd* allele number (allele-70) was assigned for the 5 VREfm isolates 111, 121, 137, 154-1, 158 by request from the database.

### Plasmid detection and typing

Each contig of <2 Mbp in size in a closed genome assembly was considered a putative plasmid. *In silico* detection and typing of plasmids was performed with PlasmidFinder (v 2.1) (23). Putative plasmids in closed genome assemblies that were not classifiable by PlasmidFinder were analyzed further (see below).

For comparison with pRUM-like/*rep17* plasmids from Dallas isolates, plasmids were extracted from PLSDB v.2021_06_23_v2 (24, 25) with the following terms: genus = *Enterococcus*, topology = circular, PlasmidFinder = contains *rep17*. Plasmids were visualized with Easyfig (26).

### Definition of pMIX plasmid groups

Putative plasmid sequences that were unclassifiable by PlasmidFinder were analyzed as follows. Presumptive *rep* genes were identified from plasmid annotations and analyzed by NCBI Conserved Domain Analysis (CDS). Confirmed *rep* genes were those that encoded protein domains that matched known Rep protein families with e-values of e^-10^ to e^-78^. The Rep sequences were also scanned by HMMER (v 2.41.1) (27) to analyze Pfam domains. Multiple sequence alignment of the nucleotide and predicted amino acid sequences of *rep* genes were performed with MUSCLE (v 3.8.424) where the “group sequences by similarity” option was selected. A neighbor-joining tree was built based on the sequence alignment using the “Tamura Nei” genetic distance model (28). The tree was visually scrutinized in order to sort the plasmids into 9 groups harboring *rep* genes with at least 95% identity in both nucleotide and predicted amino acid sequence (except for pMI4; see Results). Each unique *rep* group of plasmids was given a name of format pMIX, where the “X” is a numeric number from 1 to 9. A representative sequence from each pMIX *rep* group is in **Dataset S1B**. Representative sequences were compared to PLSDB v.2021_06_23_v2 (24, 25) using BLASTn with 90% identity and query coverage thresholds.

Eight closed circular DNA elements of varying sizes (3 to 72 kb) could not be classified by PlasmidFinder nor by the pMIX *rep* typing scheme described above. Analysis of these elements is described in the Supplemental Text.

### Average nucleotide identity (ANI) analysis

All vs. all ANI was calculated using ANIclustermap v1.1.0 at default parameters with either all *E. faecium* assemblies or *rep17*/pRUM plasmids as input.

### Transcriptional activity of predicted toxin-antitoxin (TA) systems

Transcriptional activity of the TA systems, TA_*axe-txe*_ and TA_*relE*_, was determined by RT-qPCR for the VREfm isolates 1 and 5, described in the Supplemental Text. The primer sequences used for qPCR are provided in **Dataset S1C.**

### Resistance gene identification

Acquired antimicrobial resistance genes were detected with ResFinder 3.1 (29).

### Tn*1546* analysis

The *Tn1546* nucleotide sequence described in *E. faecium* BM4147 (accession number M97297) was used as a reference to identify variations occurring in the Tn*1546* identified in this study. The Tn*1546* were classified using a previously described nomenclature (13). The presence of insertion sequence elements (IS elements) was denoted with a one-letter code, shown in parentheses: IS*1216* (B), *IS1251* (C), *ISEfa5* (D), IS*256* (J), *ISEfa16* (K), *ISEfa17* (L), *ISEfa18* (M). The combination of IS elements within the transposon is described by a two or three letter code; i.e., group ‘BC’ possesses one each of IS*1216* and IS*1251*. Arabic numerals, following the alphabet code, indicate differences due to point mutations or different IS element insertion sites. For the group ‘BC’, a previous study (13) described BC1-BC5, each with specific insertion sites of IS elements. Any novel insertion sites identified in this present study were therefore numbered from ‘BC6’ forward. The novel *ISEfa16, ISEfa17, ISEfa18* sequences were identified in this study and registered in the ISFinder public database (30).

### Accession numbers

DNA sequences from this study have been deposited under Bioproject PRJNA682584. Individual accession numbers for all sequence files are provided in **Dataset S1A**.

## Results

### ST17 and ST18 VREfm isolates were prevalent in the collection

Genome sequencing and hybrid assembly were performed for 47 *E. faecium* isolates with Oxford Nanopore MinION and Illumina technologies. 31 closed and 16 draft genome assemblies were generated (**Dataset S1A**). In our previous analysis of these isolates using PCR, all isolates were found to be *vanA*-positive and *vanB*-negative, with the exception of the isolate 163-1, which had neither *vanA* or *vanB* and had a vancomycin MIC of 2 μg/mL by broth microdilution (16). We included 163-1 in this study for comparative purposes to evaluate its relationship to the *vanA*-positive isolates in the collection. After recovery from Spectra VRE rectal surveillance cultures, all 46 *vanA*-positive isolates sequenced in this study grew on agar supplemented with 256 μg/mL vancomycin. Subsequent broth microdilution assays performed over a year later after genome sequencing and freezer restocking re-confirmed vancomycin resistance for all isolates except for 9-2 and 154-1, which had vancomycin MICs ≤ 4 μg/mL. Likely, loss of vancomycin resistance elements during laboratory culture led to these phenotypes; however, this was not investigated further in this study.

Phylogenetic analysis based on alignment of 1706 core genes allowed us to assess the diversity of the VREfm isolates (**Figure 1**). Two reference genomes, *E. faecium* 1,231,501 (5) and *E. faecium* ATCC^®^ 700221™(31), were included as representatives of *E. faecium* clade A2 and A1, respectively. *E. faecium* 1,231,501 is a human bloodstream isolate from the USA and is susceptible to vancomycin. *E. faecium* ATCC^®^ 700221 ™ is a USA isolate and VanA-type VREfm control strain used in antimicrobial susceptibility testing. As expected, the Dallas VREfm isolates clustered with the clade A1 reference *E. faecium* ATCC^®^ 700221 ™.

**Figure 1.**
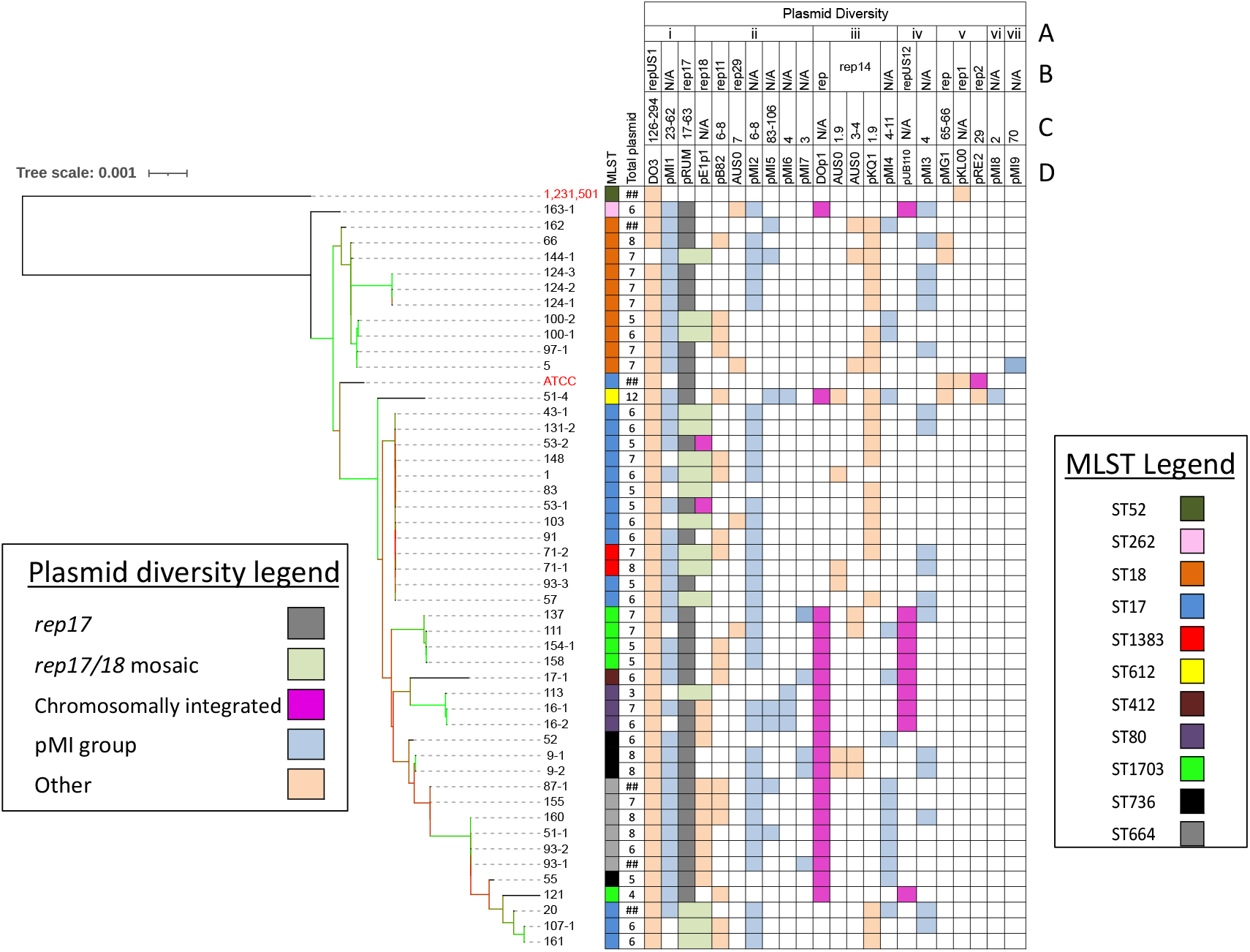
Phylogeny and plasmid content of Dallas *E. faecium* isolates. The phylogenetic tree of 49 *E. faecium* strains is based on an alignment of SNPs in 1706 core genes. The branches colored green, golden and red are supported by a bootstrap value of >70, 40-60, and <30 respectively. The reference genomes *E. faecium* 1,231,501 and ATCC^®^ 700221™ are labeled in red. Data for MLST, number of presumptive plasmids (shown as ## for incomplete assemblies), and plasmid *rep* types are shown. Plasmid replicon conserved domains i) RepA_N, ii) Rep_3, iii) Rep_trans, iv) Rep1, v) Inc18, vi) Rep_2, vii) Rol_Rep_N **(A)**, *rep* family **(B)**, plasmid size ranges **(C)**, and reference plasmid names **(D)** present in each strain are shown. A dark purple box indicates a chromosomally integrated plasmid whose size range was not determined. The *rep17* and mosaic of *rep17/rep18* are indicated with the gray and light green boxes, respectively. Plasmid groups defined in our study (pMI1 to pMI9; see main text) are highlighted in light blue color.

Ten different sequence types (ST), all belonging to clonal complex 17 (CC17), were identified in the collection (**Figure 1, Figure 2**). Isolates belonging to ST17 and ST18 are the most prevalent in the collection, comprising 50% of the isolates, which clustered together in the core genome tree (**Figure 1, Figure 2**). Six STs (ST18, ST664, ST1703, ST736, ST80 and ST612) are double locus variants of ST17. ST262 and ST412 are triple locus variants and ST1383 is a single locus variant of ST17 (**Dataset S1D**). ST1703, a double locus variant of ST17 (*pstS* allele-20; novel *gyd* allele-70), is a novel ST in the MLST database and comprised 11% of the collection. We conclude that *E. faecium* of multiple different STs, all within CC17, colonized the Dallas patients sampled at the time of the study.

**Figure 2:**
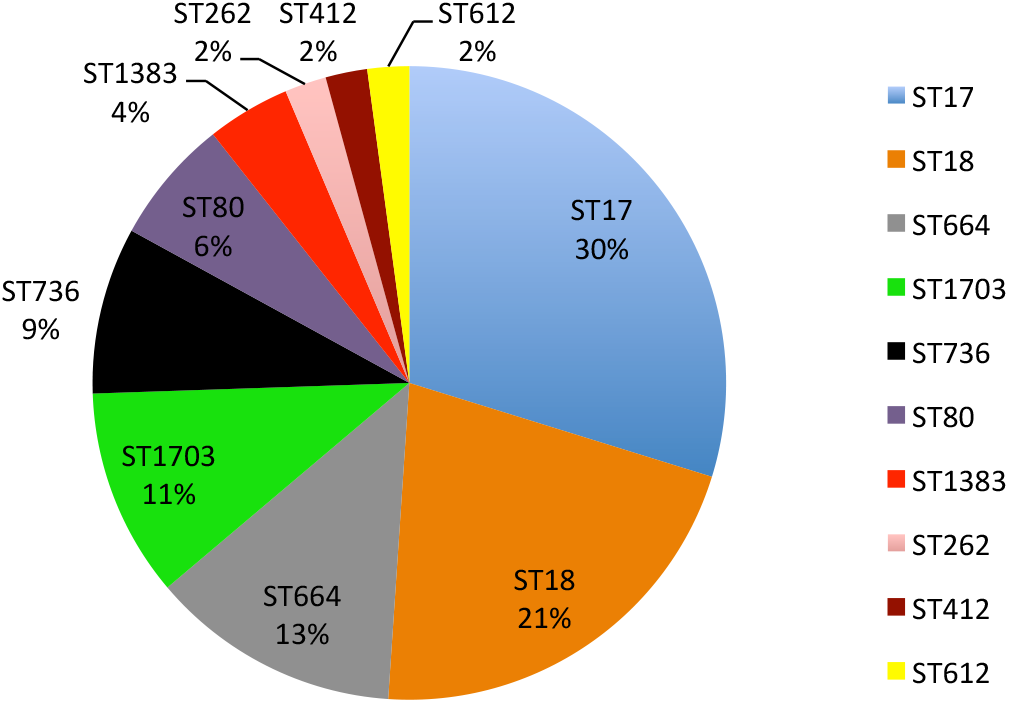
Sequence type (ST) distribution of Dallas *E. faecium* isolates. A total of 10 STs were observed across 47 isolates. The frequency of STs (percent of total isolates;) is shown.

We performed an additional analysis to determine how closely related *E. faecium* strains are in our collection. All vs. all Average Nucleotide Identity (ANI) was calculated with all *E. faecium* assemblies as input (**Dataset S1E**). Dallas *E. faecium* isolates possess ≥98.98% ANI, consistent with the core gene phylogeny in **Figure 1**. In most cases, strains isolated from the same patient were identical or nearly so. For example, the three isolates from patient 124 shared 99.99% ANI. Similar observations were made for isolates from patients 100, 16, 53, 71, and 9. Exceptions were patients 51 and 93, which each were colonized with different *E. faecium* STs (**Figure 1**), also reflected in the ANI analysis of the isolates (≤99.4% ANI).

### Large numbers of plasmids are present in the Dallas isolates

The plasmid content of the Dallas isolates was analyzed by an *in silico* detection method utilizing the conserved nucleotide sequences of replication initiation (*rep*) genes (23). Each contig of <2 Mbp in size in a closed genome assembly was considered a putative plasmid. Of the 259 closed circular putative plasmids, 141 were assigned to 10 known *rep* families described in the PlasmidFinder database. The remaining 118 putative plasmids were categorized into 9 novel plasmid groups (pMI1 to pMI9) and 8 other elements (discussed in the next section).

The Dallas isolates harbored a variable number of putative plasmids (n=3 to 12) of sizes ranging from 1.9 to 294 kb (**Figure 1; Dataset S1F**). The most common elements were a megaplasmid (126 to 294 kb) and a pRUM-like plasmid (17 to 63 kb), which were each present in all isolates except 144-1, which lacked the megaplasmid. Both the reference strains (*E. faecium* 1,231,501 and ATCC^®^ 700221™) harbored the megaplasmid, but 1,231,501 lacked the pRUM-like plasmid. Each of the megaplasmids harbored the *rep* gene *repUS15*, encoding a RepA_N protein motif. The pRUM-like plasmid harbored *rep17* or both *rep17* and *rep18*, which we refer to as a mosaic plasmid. An analysis of plasmid replicon enrichment among specific STs is presented in the **Supplemental Text** and **Dataset S1G**.

### Uncategorized plasmids were sorted into 9 plasmid groups, pMI1 to pMI9

PlasmidFinder (23) did not identify *rep* genes in a total of 118 circular entities. Of these 118 circular entities, 110 were established to be plasmids based on the presence of *rep* genes not currently represented in the PlasmidFinder database (see Materials and Methods). The remaining 8 circular entities were designated as putative excision products of genomic islands (**Supplementary Text, Dataset S1H-I**) including a 5 kb circular fragment of Tn*1546* in isolate 111.

The 110 plasmids not classifiable by PlasmidFinder were sorted into 9 plasmid groups, pMI1 to pMI9, based on phylogenetic analyses of the *rep* genes (**Figure S1; Dataset S1J**). The nucleotide sequences of the *rep* genes and their corresponding amino acid sequences shared >95% identity within each pMIX plasmid group, except for pMI4 (**Table 1**). The clustering of the *rep* gene sequences belonging to the pMI4 plasmid group supported their inclusion in a single group, despite lower pairwise sequence identities (**Figure S1**). A total of 6 Rep protein families (RepA_N, Rep_3, Rep_1, Rep_trans, Rol_rep_N, and Rep_2) were encoded by the *rep* genes of the 9 pMIX groups (**Dataset S1J**). Specifically, the *rep* genes of pMI1 and pMI3 encoded protein families RepA_N and Rep_1, respectively. The *rep* genes of pMI2, pMI5, pMI6 and pMI7 encoded the protein family Rep_3. The *rep* genes of pMI4 and pMI8 encoded protein families Rep_trans and Rep_2, respectively. The *rep* of pMI9 was the only exception, encoding two Rep protein families, Rep_trans and Rol_rep_N (**Table 1**).

**Table 1.**
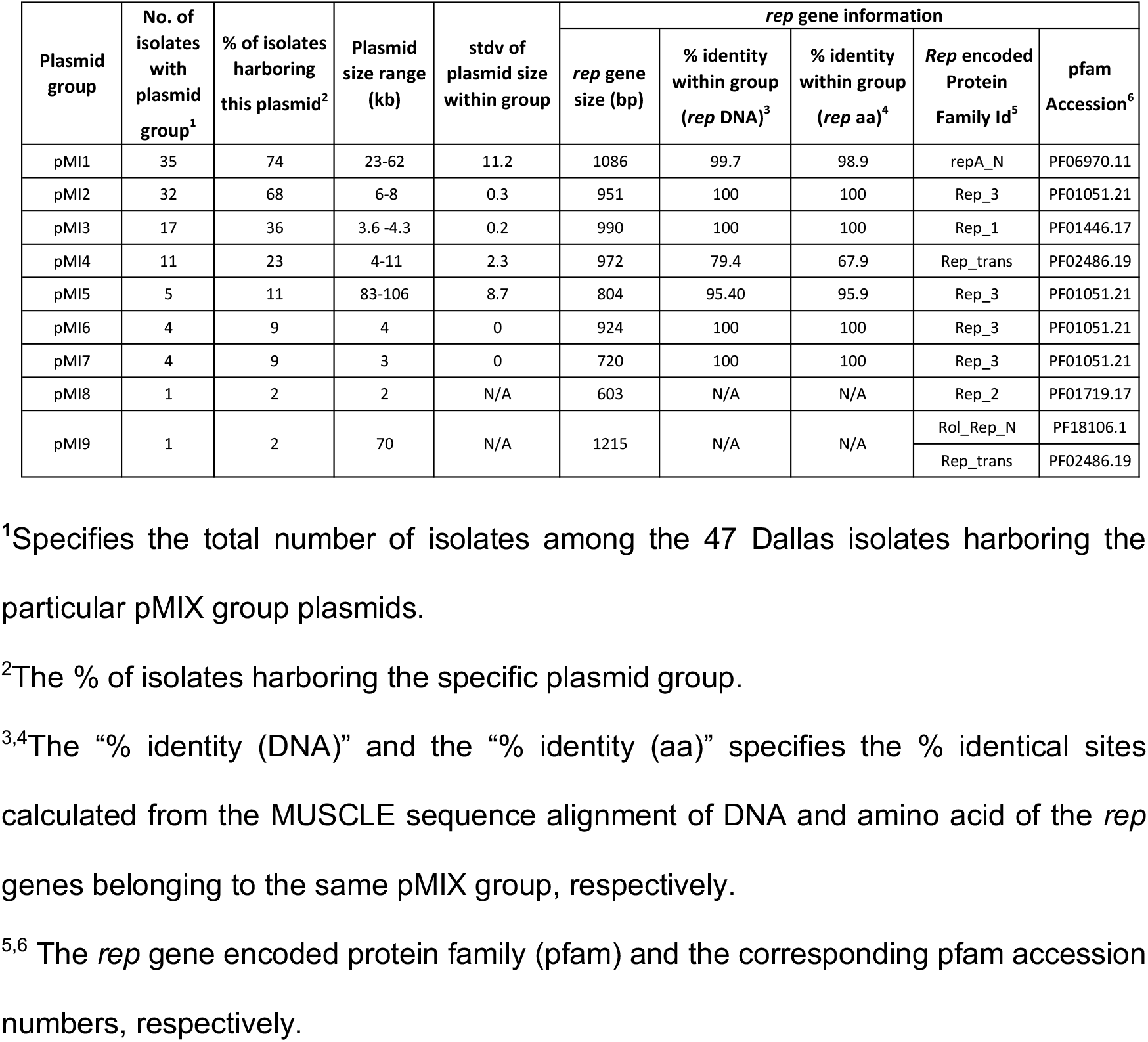
Features of the pMIX group plasmids.

Each of the Dallas isolates harbored 1 to 5 of the pMIX plasmid groups, except the isolate 83 (ST17), which had none of them (**Figure 1**). Neither of the two reference strains harbored any of the pMIX plasmid groups. The pMI1 was the most frequently occurring pMIX plasmid group, identified in 75% of the isolates (**Table 1**). The pMI8 and pMI9 were the rarest ones, found in only isolates 51_1 (ST612) and 5 (ST18), respectively. The pMIX plasmid groups were not enriched in any specific ST. We observed that plasmids within a very narrow size range (standard deviation 0.2 to 2.3) were grouped together by this *rep* typing scheme (with the exception of pMI1), although plasmid size was not taken into consideration in assigning plasmid groups.

We used PLSDB (24, 25) to compare representative *rep* sequences from the pMIX plasmid groups with existing plasmids in the database (as of January 2023). PLSDB compiles plasmid sequences and their associated metadata, including PlasmidFinder and other analyses. This was done as an additional check on our analysis, as ours was initially performed in 2018-2019. We reasoned that the pMIX typing scheme would be most useful if it encompassed other previously untypeable plasmids in the database. We applied strict thresholds of 90% identity and 90% query coverage to filter BLASTn hits to the database. The results support pMI1, pMI3, pMI5, pMI6, and pMI8 as newly defined *rep* types that include other previously untyped *Enterococcus* plasmids from around the world (**Dataset S1K**). The only hit to pMI9 *rep* was the Dallas isolate from which it was defined. In this PLSDB analysis, pMI2 was assigned to *rep11a*, and pMI4 was assigned to *rep14b*. The pMI7 analysis was more complicated, with one group of identical hits (100% identity and query coverage) for many plasmids with no known *rep* type assigned, and another group of hits with 100% identity but lower 98% coverage that were assigned to *rep17*.

### Resistance genes were encoded in both chromosomes and plasmids

Different combinations of 12 resistance genes predicted to provide resistance against 5 classes of antibiotics (aminoglycosides, glycopeptides, macrolides, tetracycline, and trimethoprim) were identified among the Dallas isolates (**Figure 3**; **Dataset S1L; Supplemental Text**). The most commonly occurring resistance genes were *aac(6’)I-1* (100% of isolates), *msr* (100%), *vanA* (95%), *tetM* (95%), *ermB* (85%), and *aph(3’*) (85%). These resistance genes were encoded in the chromosome and primarily in only two plasmid types, the pRUM-like plasmid group and the megaplasmid. The pRUM-like plasmids predominantly encoded 4 of the resistance genes (*ant(6)-Ia, aph(3’)III, vanA*, and *ermB*) conferring resistance against aminoglycosides, vancomycin and erythromycin (**Figure 3**). The *repUS7* plasmid (pMG1) was found among only 3 of the Dallas isolates - 66 (ST18), 144-1 (ST18) and 51-4 (ST612) - and encoded *dfrG*, for trimethoprim resistance. We note that antibiotic resistance of these isolates was not phenotypically assessed in this study, other than for vancomycin resistance.

**Figure 3.**
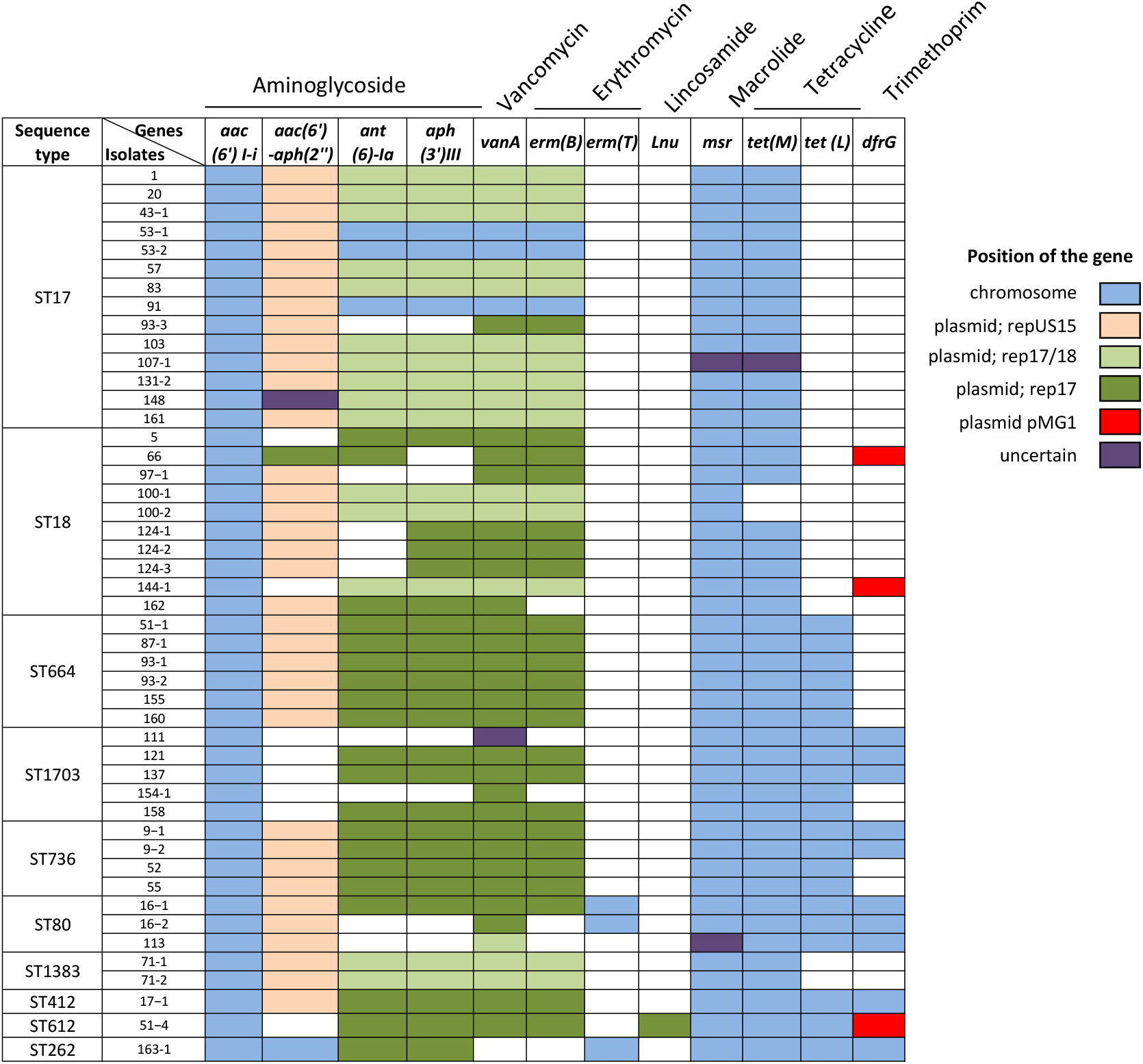
Distribution of predicted antibiotic resistance genes among Dallas isolate genomes. Shown are 12 resistance genes (6’-N-aminoglycoside acetyltransferase, (*aac(6’)I-i*), bifunctional enzyme 6’-aminoglycoside acetyltransferase-2”-aminoglycoside phosphotransferase (*aac(6’)-aph(2”)*), aminoglycoside nucleotidyltransferase (*ant(6)-Ia), aph(3’)III, vanA, erm(B), erm(T), Inu*, acquired macrolide resistance-like protein (*msr), tet(M), tet(L) and* dihydrofolate reductase (*dfrG)*), and the antibiotics they provide resistance to. A solid colored box indicates the presence of a certain resistance gene in the isolate. The boxes are color coded to indicate the location of the gene (chromosome or plasmid).

All of the Dallas isolates, except for 163-1, encoded VanA-type vancomycin resistance genes. The *vanA* genes possessed 99.9% nucleotide and 99.7% amino acid sequence identity with the reference *vanA* sequence (GenBank AAA65956.1 (**Dataset S1L).** The reference sequence is encoded within the transposon Tn*1546* (GenBank M97297), originally described in the human fecal isolate *E. faecium* BM4147 collected from France in 1988 (32) (**Figure S2**). A classical structure of the *vanA* gene cluster, consisting of *vanHAX* and *vanYZ* encoded within Tn*1546* elements, was observed among the Dallas VREfm isolates, except for the isolate 17-1, which lacked the genes *vanY* and *vanZ*. The Tn*1546* structure varied among the isolates in terms of insertion site (chromosome versus plasmid), IS elements, and point mutations, discussed further below.

### The Tn*1546* is borne on pRUM-like plasmids in most isolates

The complete Tn*1546* was carried by the pRUM-like plasmid in 42 of 46 *vanA*-positive isolates. A representative pRUM-like plasmid from our collection is shown in **Figure 4**. The isolate 111 did not encode the entire Tn*1546* in the pRUM-like plasmid; rather, it encoded *vanRS* in a 17 kb pRUM-like plasmid while *vanHAX* was encoded within a separate 5 kb circular element (te111_5kb) (**Dataset S1H**). Tn*1546* was chromosomally encoded in 3 isolates, 53-1, 53-2 and 91, each belonging to ST17, with all three strains also harboring a pRUM-like plasmid lacking *Tn1546* (**Table 2**).

**Figure 4.**
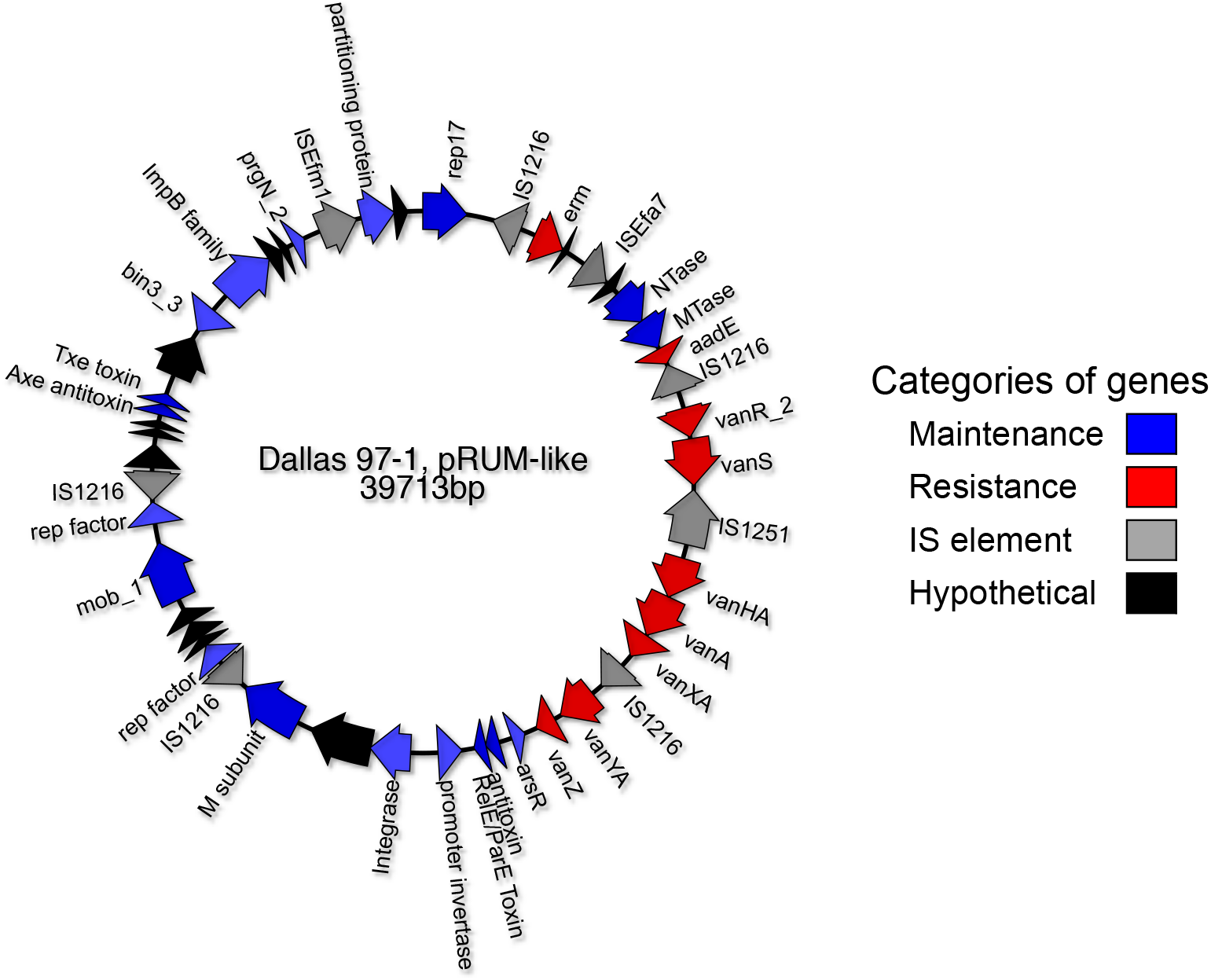
Genetic map of a representative Group 2a pRUM-like plasmid. The 39 kb plasmid p97_1_39kb (accession number CP066571) from the Dallas VREfm isolate 97-1 is shown. Arrows represent genes. The genes are color-coded based on their predicted activity. Maintenance genes are predicted to be responsible for general maintenance of the plasmid within the bacterial host. Resistance genes provide antibiotic resistance.

**Table 2.**
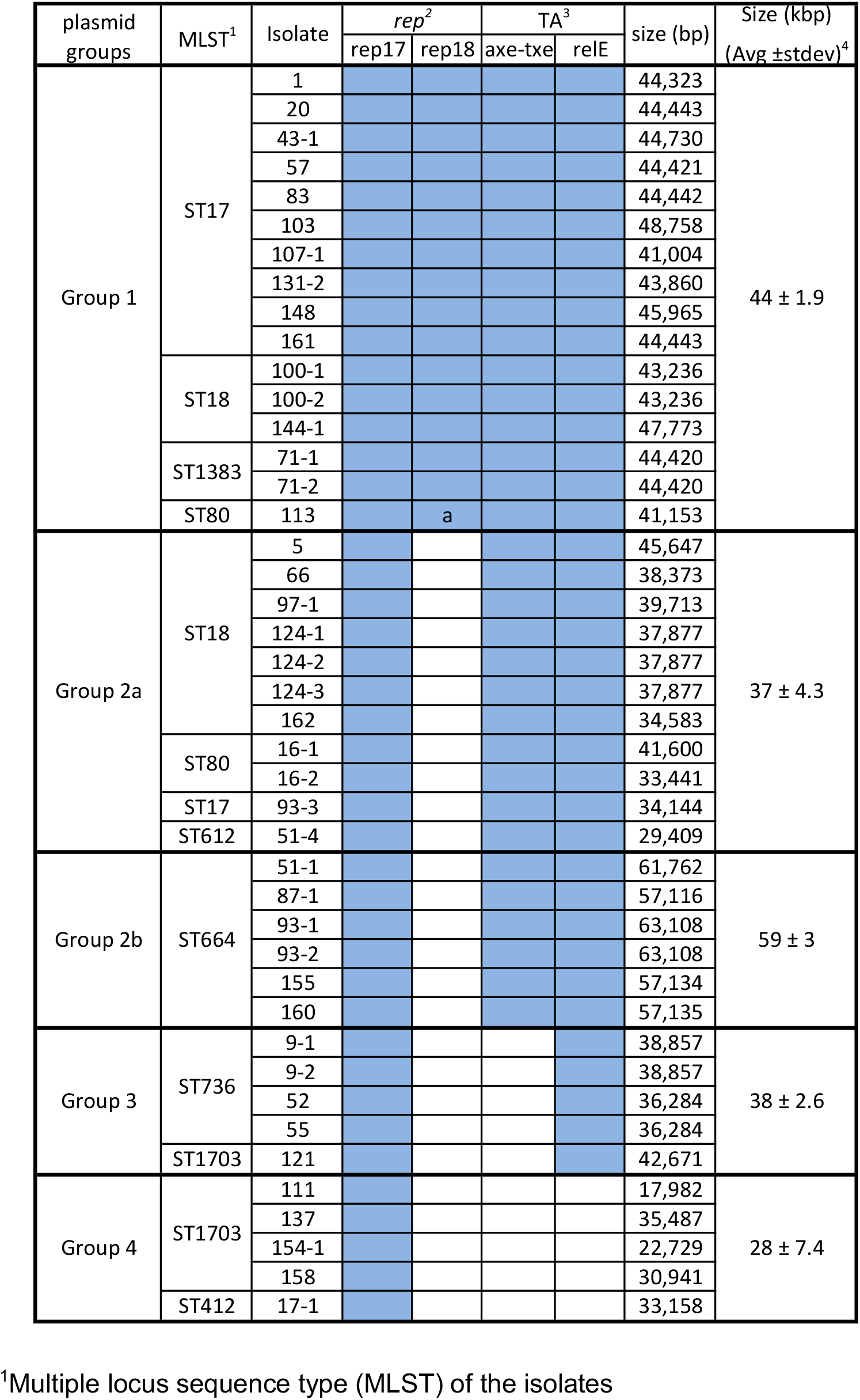
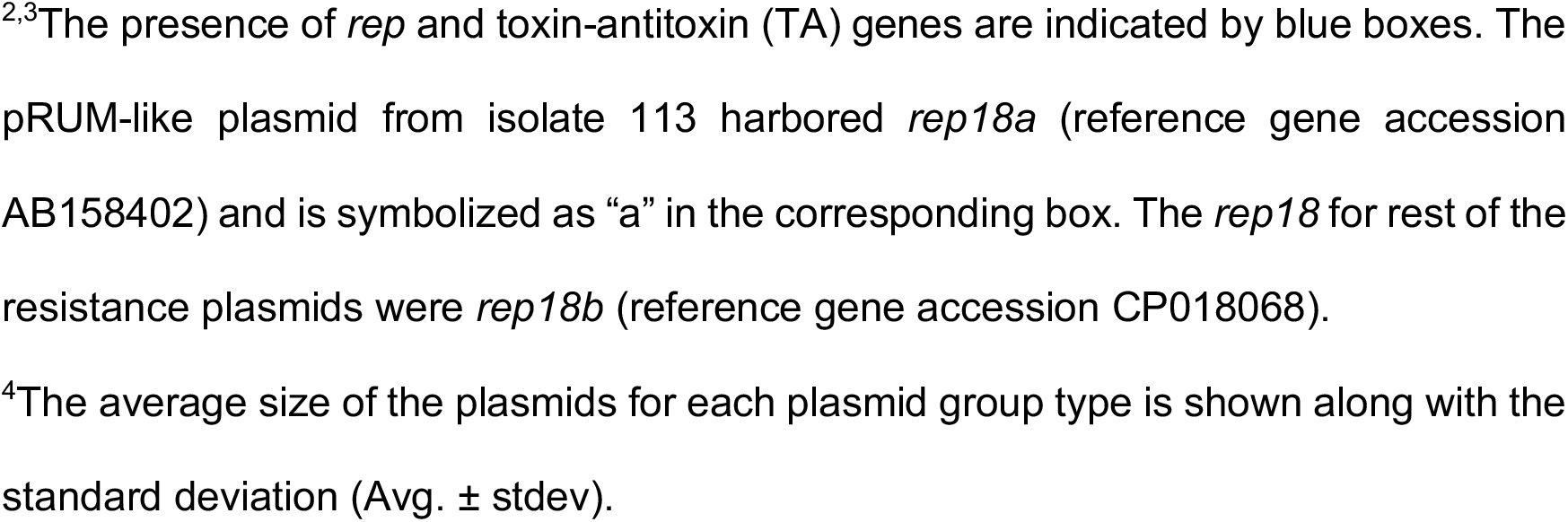
Features of the pRUM-like plasmid groups.

| plasmid groups | MLST<sup>1</sup> | Isolate | *rep*<sup>2</sup> |  | TA<sup>3</sup> |  | size (bp) | Size (kbp)<br>(Avg ±stdev)<sup>4</sup> |
| --- | --- | --- | --- | --- | --- | --- | --- | --- |
|  |  |  | rep17 | rep18 | axe-txe | relE |  |  |
| Group 1 | ST17 | 1 |  |  |  |  | 44,323 | 44 ± 1.9 |
|  |  | 20 |  |  |  |  | 44,443 |  |
|  |  | 43-1 |  |  |  |  | 44,730 |  |
|  |  | 57 |  |  |  |  | 44,421 |  |
|  |  | 83 |  |  |  |  | 44,442 |  |
|  |  | 103 |  |  |  |  | 48,758 |  |
|  |  | 107-1 |  |  |  |  | 41,004 |  |
|  |  | 131-2 |  |  |  |  | 43,860 |  |
|  |  | 148 |  |  |  |  | 45,965 |  |
|  |  | 161 |  |  |  |  | 44,443 |  |
|  | ST18 | 100-1 |  |  |  |  | 43,236 |  |
|  |  | 100-2 |  |  |  |  | 43,236 |  |
|  |  | 144-1 |  |  |  |  | 47,773 |  |
|  | ST1383 | 71-1 |  |  |  |  | 44,420 |  |
|  |  | 71-2 |  |  |  |  | 44,420 |  |
|  | ST80 | 113 |  | a |  |  | 41,153 |  |
| Group 2a | ST18 | 5 |  |  |  |  | 45,647 | 37 ± 4.3 |
|  |  | 66 |  |  |  |  | 38,373 |  |
|  |  | 97-1 |  |  |  |  | 39,713 |  |
|  |  | 124-1 |  |  |  |  | 37,877 |  |
|  |  | 124-2 |  |  |  |  | 37,877 |  |
|  |  | 124-3 |  |  |  |  | 37,877 |  |
|  |  | 162 |  |  |  |  | 34,583 |  |
|  | ST80 | 16-1 |  |  |  |  | 41,600 |  |
|  |  | 16-2 |  |  |  |  | 33,441 |  |
|  | ST17 | 93-3 |  |  |  |  | 34,144 |  |
|  | ST612 | 51-4 |  |  |  |  | 29,409 |  |
| Group 2b | ST664 | 51-1 |  |  |  |  | 61,762 | 59 ± 3 |
|  |  | 87-1 |  |  |  |  | 57,116 |  |
|  |  | 93-1 |  |  |  |  | 63,108 |  |
|  |  | 93-2 |  |  |  |  | 63,108 |  |
|  |  | 155 |  |  |  |  | 57,134 |  |
|  |  | 160 |  |  |  |  | 57,135 |  |
| Group 3 | ST736 | 9-1 |  |  |  |  | 38,857 | 38 ± 2.6 |
|  |  | 9-2 |  |  |  |  | 38,857 |  |
|  |  | 52 |  |  |  |  | 36,284 |  |
|  |  | 55 |  |  |  |  | 36,284 |  |
|  | ST1703 | 121 |  |  |  |  | 42,671 |  |
| Group 4 | ST1703 | 111 |  |  |  |  | 17,982 | 28 ± 7.4 |
|  |  | 137 |  |  |  |  | 35,487 |  |
|  |  | 154-1 |  |  |  |  | 22,729 |  |
|  |  | 158 |  |  |  |  | 30,941 |  |
|  | ST412 | 17-1 |  |  |  |  | 33,158 |  |

<sup>1</sup>Multiple locus sequence type (MLST) of the isolates
<sup>2,3</sup>The presence of *rep* and toxin-antitoxin (TA) genes are indicated by blue boxes. The pRUM-like plasmid from isolate 113 harbored *rep18a* (reference gene accession AB158402) and is symbolized as “a” in the corresponding box. The *rep18* for rest of the resistance plasmids were *rep18b* (reference gene accession CP018068).
<sup>4</sup>The average size of the plasmids for each plasmid group type is shown along with the standard deviation (Avg. ± stdev).

We categorized the 43 pRUM-like plasmids encoding Tn*1546* or a fragment of Tn*1546* (for isolate 111) into four groups based on the presence or absence of replication and stability modules on their backbones (**Table 2, Figure 5**), per a previously described typing scheme (14). For the plasmids in our collection, two toxin-antitoxin (TA) systems were observed in different distributions, *axe-txe* (TA_axe-txe_) and a *relE/parE* family toxin associated with an uncharacterized antitoxin component (TA_relE_) (**Table 2**). Both of the TA systems were actively expressed in two representative VREfm isolates (**Figure S3**).

**Figure 5.**
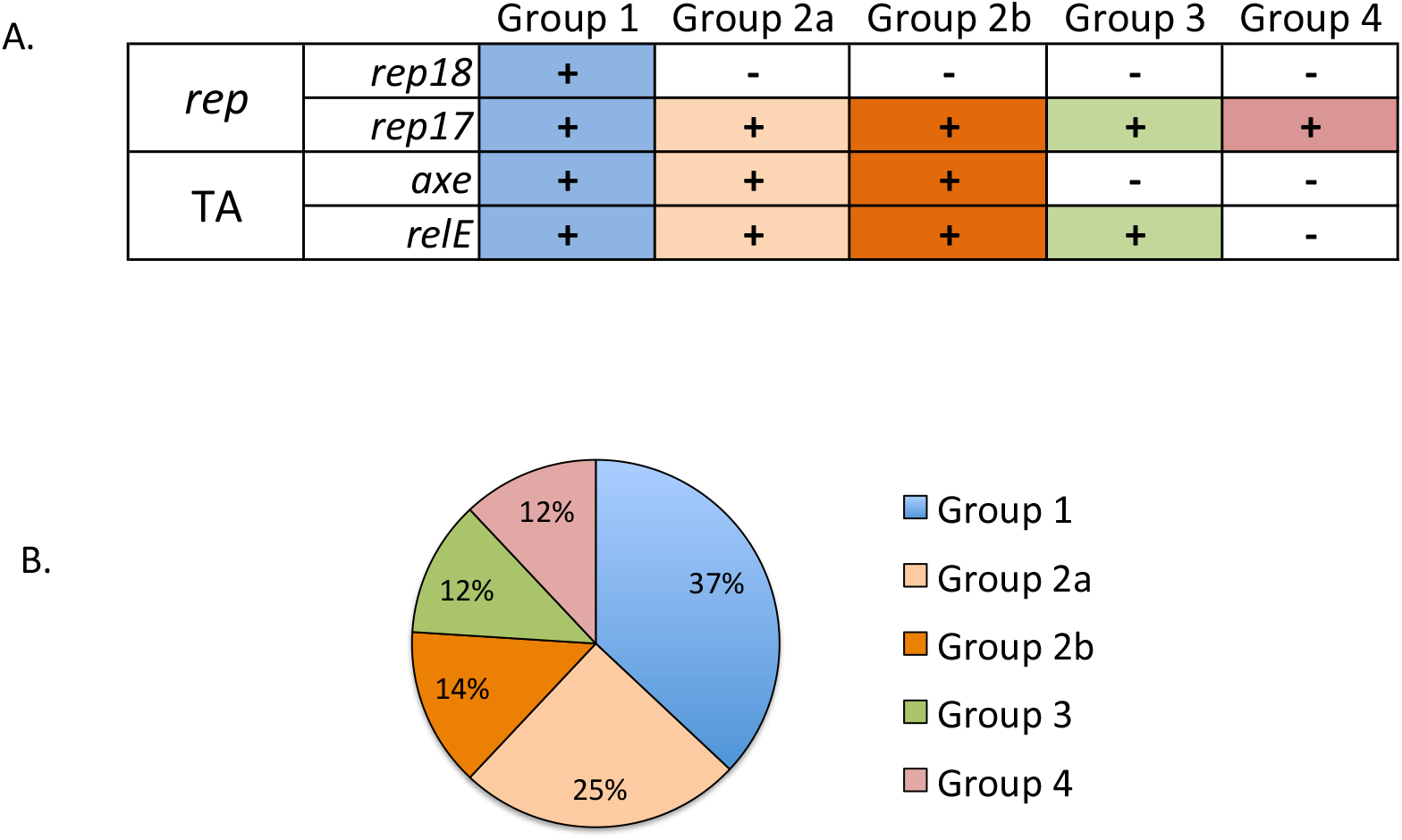
Classification and distribution of pRUM-like plasmid groups. **A**) pRUM-like plasmids were classified based on the presence or absence (indicated by a + or -, respectively) of 2 replication modules (*rep17*and *rep18*) and 2 stability modules (toxin-antitoxin system; TA_axe-txe_ and TA_relE_). **B)** Frequency of the pRUM-like plasmid groups among 43 Dallas isolates.

i. Group 1 (rep_17_; rep_18_; TA_axe-txe_; TA_relE_; n=16): These plasmids possess *rep_17/18_* and both TA_axe-txe_ and TA_relE_. All the group 1 plasmids encoded the *rep18b* (accession number CP018068), except isolate 113, which encoded the *rep18a* (accession number AB158402). The plasmids had a narrow size range (44 to 48 kb) and were present in VREfm isolates from ST17, ST18, ST1383 and ST80.
ii. Group 2 (rep_17_; TA_axe-txe_; TA_relE_; n=17): These plasmids possess *rep_17_* and both TA_axe-txe_ and TA_relE_. The group 2 plasmids were classified further based on size and genetic background of the isolates carrying them. Group 2a plasmids (29-45 kb) were carried by the reference strain ATCC^®^ 700221 ™, and isolates belonging to ST18 (5, 66, 97-1, 124-1, 124-2, 124-3, and 162), ST80 (16-1, 16-2), ST17 (93-2), and ST612 (51-4). The comparatively larger (57-63 kb) group 2b plasmids were present in all 6 isolates belonging to ST664 (51-1, 87-1, 93-1, 93-2, 155, and 160). Gene presence/absence analysis using Roary (19) identified a set of genes that are specific to Group 2b plasmids (**Dataset S1M**), including a ~7.5 kb region encoding helix-turn-helix XRE-family like proteins (cd00093) inserted immediately upstream of the *van* gene cluster (**Figure S4**).
iii. Group 3 (rep_17_; TA_relE_; n=5): These plasmids possess *rep_17_* and TA_relE_, and have a size range of 36-42 kb. They were carried by ST736 isolates (isolates 9-1,9-2, 52, and 55) and one ST1703 isolate (isolate 121).
iv. Group 4 (rep_17_; n=5): These plasmids possess *rep_17_* and no known stability modules, and have a size range of 17-35 kb. These plasmids were carried by isolates belonging to the novel genetic background ST1703 (111, 137, 154-1, and 158) and by a single ST412 isolate (17–1).

We further analyzed the pRUM-like plasmids in our collection using two approaches: a core genome phylogeny based on sequence variation in 7 pRUM-like plasmid core genes (*rep17, IS1216, vanS*, ImpB/MucB/SamB family protein, hypothetical protein, plasmid partitioning protein, and replication control protein PrgN) (**Figure S5**), and an analysis of ANI **(Figure S6; Dataset S1N).** The pRUM-like plasmids form two clusters in the phylogenetic tree (**Figure S5)**, which correspond to the presence/absence of TA_*axe-txe*_. The pRUM-like plasmids ranged from 93.9 – 100% pairwise ANI. ANI analysis identified well-defined clusters of plasmids with high percent ANI that correspond to plasmid groups 2b, 3, and 4, while groups 1 and 2a are intermixed (**Figure S6**). This is also consistent with the presence/absence of TA_*axe-txe*_ being an important discriminator for grouping of pRUM-like plasmids.

Finally, we compared pRUM-like/*rep17* plasmids from the Dallas collection with other *rep17* plasmids that have previously been published. PLSDB was queried to obtain circular plasmids assigned to genus *Enterococcus* and to *rep17*. A strict cut-off of ≥99.9% pairwise ANI was applied, which identified a cluster of closely related pRUM-like/*rep17* plasmids from different sources (**Figure 6**). These plasmids have ≥99.9% ANI, are from geographically distinct regions (USA [this study], France (33), India (34), and Australia (35, 36)), and are present in different *E. faecium* STs. The most closely related are the pRUM-like/*rep17* plasmid from Dallas isolate 16-1 and the *rep17* plasmid from the VREfm strain 15-307.1. These plasmids have identical gene content and differ by SNPs and deletions/insertions at 10 sites. VREfm 15-307.1 was isolated in France in 2015 from a rectal swab of a hospitalized patient, which is similar to the collection of Dallas 16-1 in 2015.

**Figure 6.**
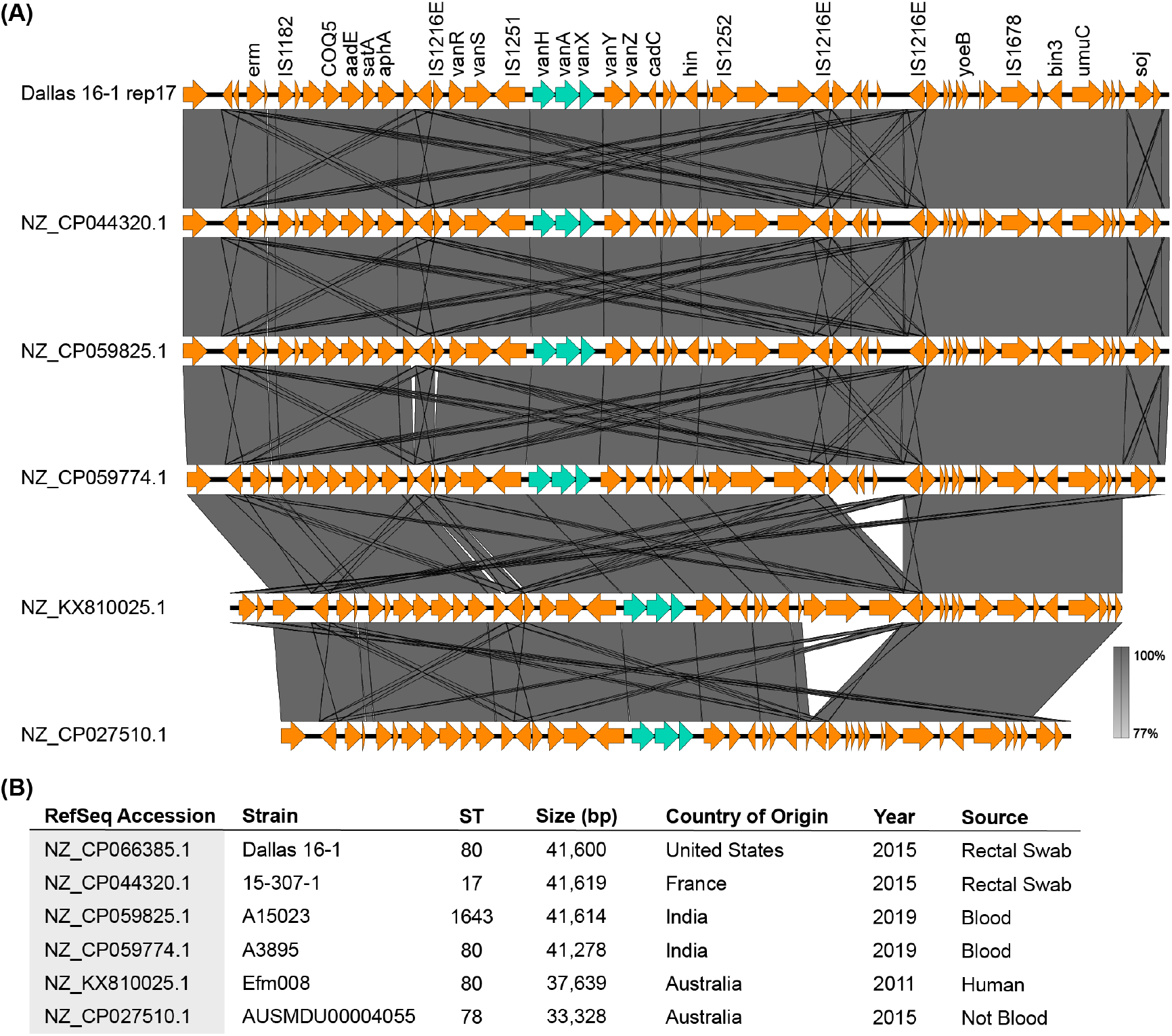
A cluster of pRUM-like plasmids identified from PLSDB with 99.9% ANI. (A) BLASTn alignment of plasmid sequences with Easyfig. (B) Data for *E. faecium* strains and plasmids, arranged in the same order as the alignment.

### Dallas VREfm harbored 17 variants of Tn*1546*

The Tn*1546* region of the Dallas isolates varied from the typical structure, which consists of ORF1 (transposase), ORF2 (resolvase) and the *van* genes flanked by IR_L_ (inverted repeat, left) and IR_R_ (inverted repeat, right) (**Figure 7**). A total of 17 different sequence variations in the Tn*1546* were observed among 45 of the Dallas isolates (**Figure 7, Table 3**). These 17 variations were classified using a nomenclature system developed previously (13) based on the presence of IS elements, point mutations, and deletions relative to the reference Tn*1546*. VREfm belonging to the same Tn*1546* group shared 99.8%-100% nucleotide sequence identity in the Tn*1546* region. The isolate 111 was excluded from this grouping since it harbored a fragmented Tn*1546* occurring in a pRUM-like plasmid and another circular entity (**Dataset S1 H**). The Tn*1546* variation was mostly associated with the distribution of 7 different IS elements (IS*1251*, IS*1216*, ISEfa*16*, *ISEfa5, ISEfa17*, IS*256*, *ISEfa18*), three of which were novel to the Dallas isolates. Most of the Tn*1546* groups described in this study are novel relative to those previously reported in the literature, except for group BC2, which was reported previously in Poland (13). The most predominant Tn*1546* groups were BC9 (n=14) and J (n=6). The BC9 variety was observed in ST17 (n=9) and ST18 (n=5) isolates. The reference strain ATCC^®^ 700221™ harbored the C2 structural variant of Tn*1546* that was described previously (13) but is not present in the Dallas VREfm isolates. The presence of different pRUM groups and Tn*1546* types across the phylogeny of *E. faecium* isolates in this study is presented in **Figure S7**.

**Figure 7:**
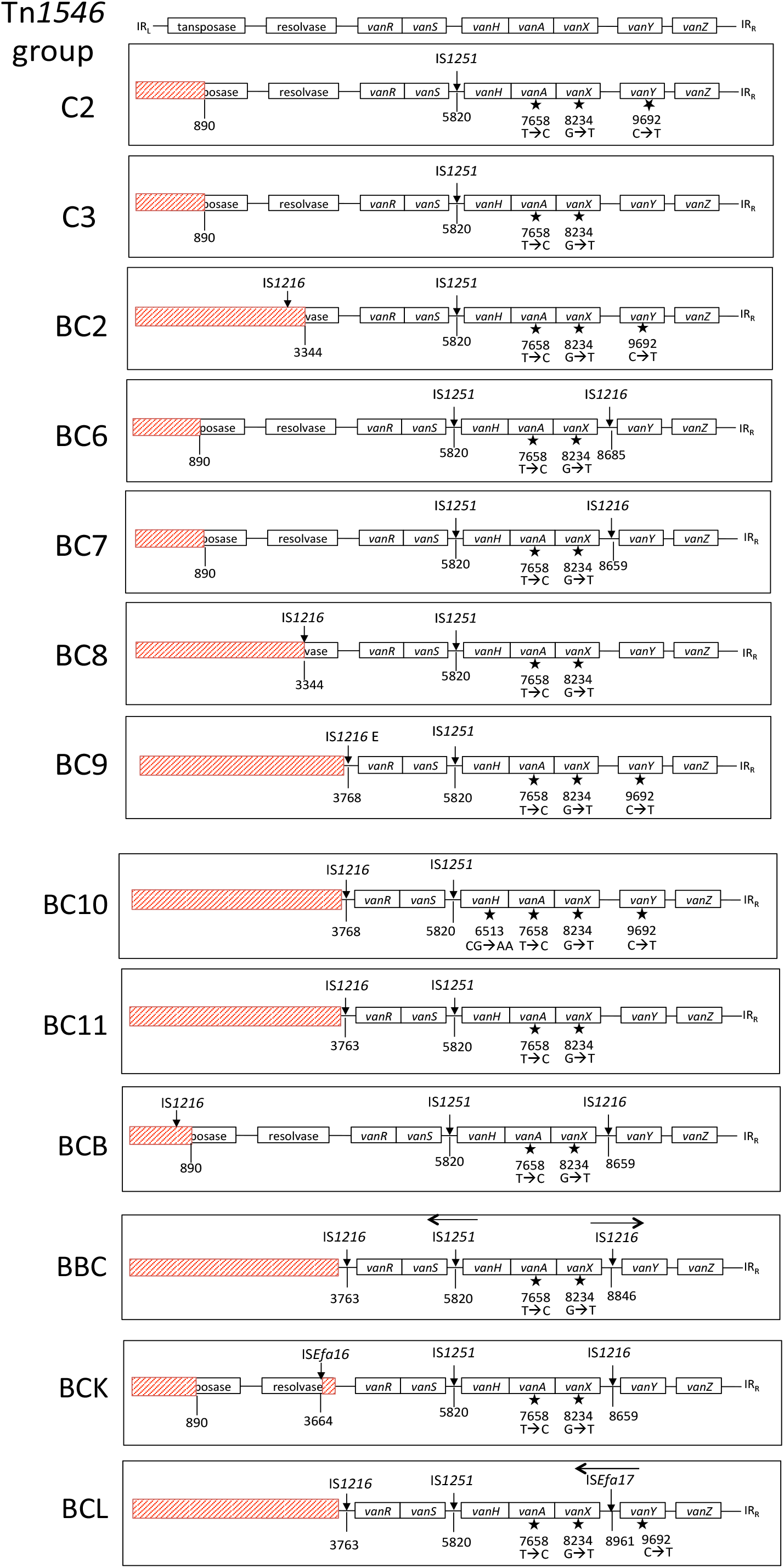
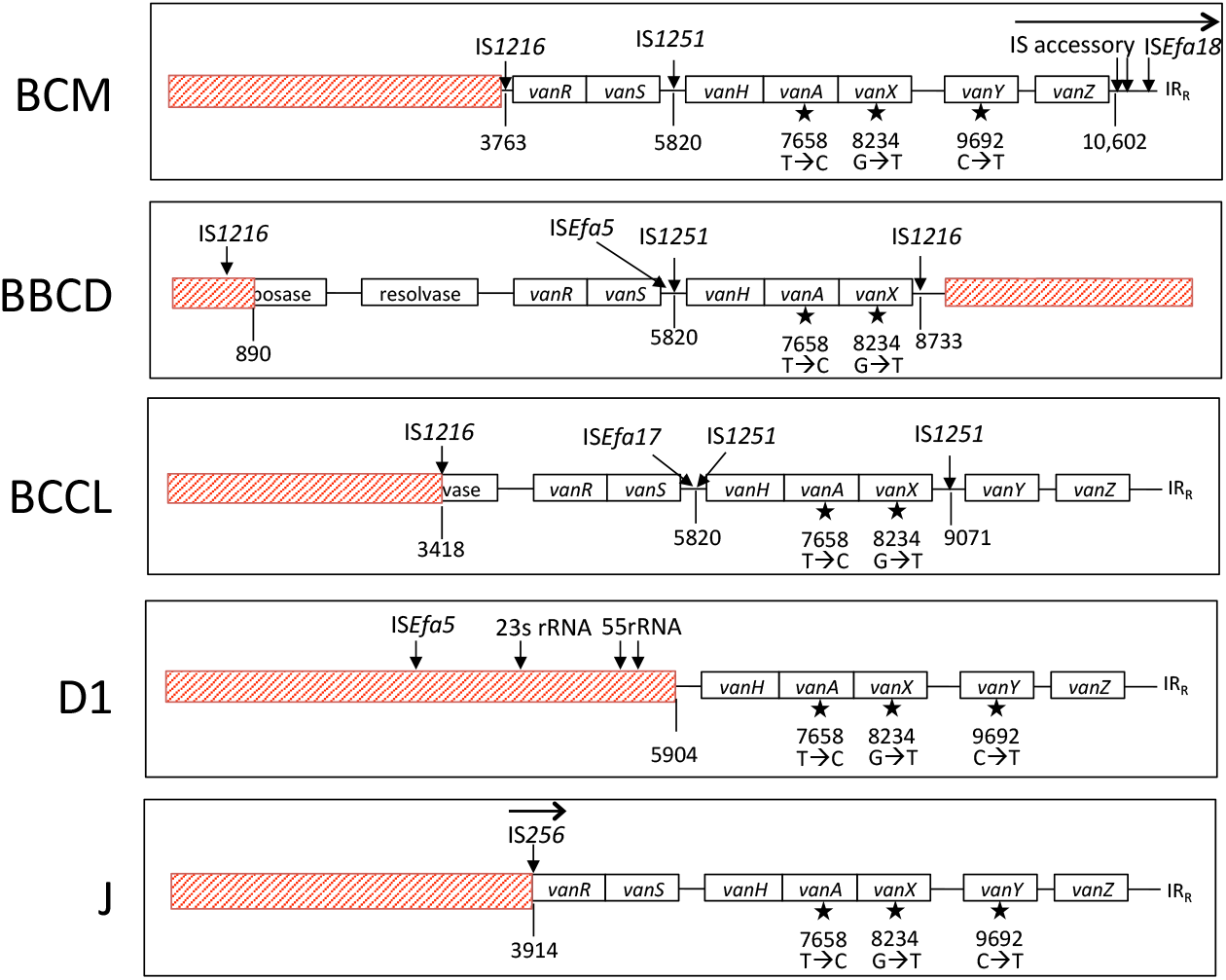
Tn*1546* structural variations in the Dallas VREfm isolates. A reference *Tn1546* structure that includes transposase, resolvase and *van* gene cluster flanked by IR_L_ and IR_R_ (inverted repeat left and right, respectively) is delineated at the top. The Tn*1546* groups are represented below the reference Tn*1546*, and the name of the group (C3, BC2, BC6, BC7, BC8, BC9, BC10, BC11, BCB, BBC, BCK, BCL, BCM, BBCD, BCCL D1, J) is shown on the left. Red boxes represent deleted regions relative to the reference Tn*1546*. Vertical lines with numbers at the bottom represent the nucleotide position within the reference *Tn1546*. IS elements are represented by downward arrows. Black stars indicate point mutations. The figure is not drawn to scale.

**Table 3.**
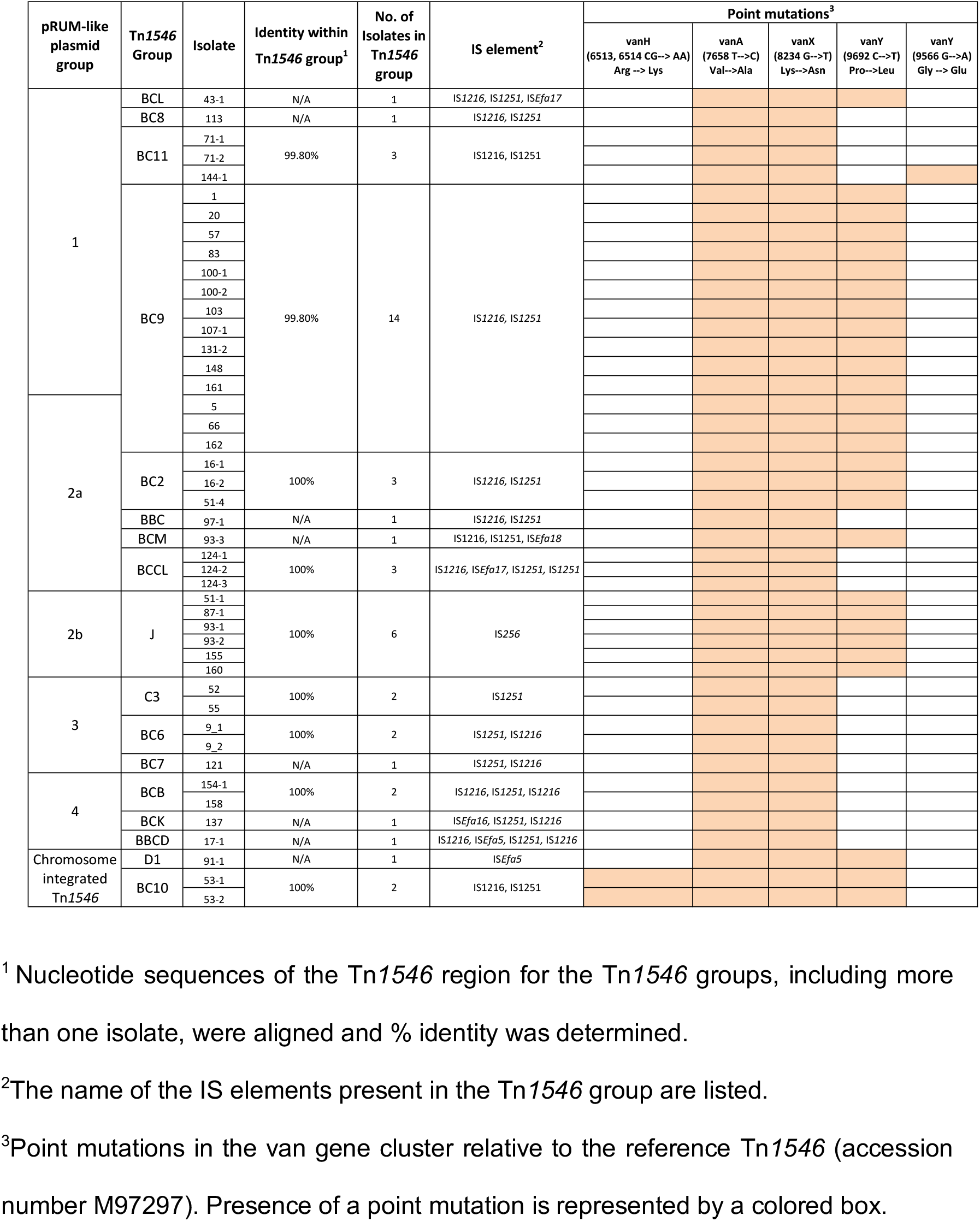
Features of the *Tn1546* groups observed among the Dallas isolates.

Specific structural variations of the Dallas Tn*1546* elements are described further in the Supplemental Text.

### Potential VREfm patient-patient transmission

We used our genome data to look for identical *E. faecium* clones colonizing different patients. This would be suggestive of prior transmission of VREfm between patients in our area. We found one possible instance supported by our genomic data. *E. faecium* isolates from patients 55 and 52 were of identical ST, same pRUM plasmid size and group, same Tn*1546* group (and are the only members of the group in this study), and possess 99.99% ANI overall. Their pRUM plasmids possess >99.99% ANI, differing by one SNP. However, the total genome size for these two strains differs. The isolate from patient 55 has a closed genome that is 1026 bp longer than the draft genome of the patient 52 isolate.

## Discussion

This study provided a snapshot of fecal VREfm colonizing hospitalized patients in Dallas, Texas from August – October 2015. VREfm isolates from previously described and new STs were detected. The most prevalent VREfm isolates, comprising 50% of those analyzed, belonged to ST17 and ST18. These STs were previously reported to be responsible for outbreaks in countries including Portugal (14), Spain (37), Ireland (38), Denmark (39), Columbia (40), Brazil (41), Canada (42). The other 8 STs identified among our collection (ST1383, ST612, ST412, ST80, ST1703 and ST664), were single, double, or triple locus variants of ST17. ST1703 is a novel sequence type described in this study. ST182, which caused an outbreak in San Antonio, Texas in the 1990s (43), was not detected in our collection.

All the plasmid-borne Tn*1546* in Dallas isolates were encoded within pRUM-like plasmids that were clustered into 4 subgroups (1, 2a, 2b, 3, and 4), defined by the presence of the replication module *rep17* and a combination of other replication and stability modules. The patients from whom the isolates in this study were collected resided in different areas of Dallas and in unconnected facilities including nursing homes and personal residences. Therefore, the presence of only one type of plasmid backbone carrying *vanA* among these patients was striking. Our data are consistent with another recent analysis of VREfm from the United States. Chilambi, et al (2020) (44) analyzed gastrointestinal and blood VREfm isolated from pediatric patients at St. Jude’s Children’s Research Hospital over a 10 year period. VanA-type VREfm was isolated from 23 of 24 patients analyzed, and for all 23, the Tn*1546* was encoded on a *rep17* element. Clearly, these plasmids are significant vectors for vancomycin resistance in *E. faecium*, as proposed by Freitas, et al. (10) in a multilevel analysis of VREfm. In our study, multiple predicted antibiotic resistance genes were detected on these plasmids, in addition to vancomycin resistance genes. The pRUM-like plasmids should be a focus of plasmid surveillance. Resistance genes may be consolidating on this specific backbone, as opposed to the many other plasmids detected in VREfm in this study.

We detected carriage of up to 12 extrachromosomal elements in the VREfm isolates in our study. This is consistent with the comprehensive work of Arredondo-Alonso, et al., who conducted a large-scale analysis of the *E. faecium* plasmidome (45). The authors determined that *E. faecium* from hospitalized patients possess the largest plasmidome in terms of plasmid number and total bp. The striking number of extrachromosomal elements among VREfm warrants further investigation into their maintenance, transmission, and functions.

## Acknowledgements

This project was supported by R01AI116610 and the Cecil H. and Ida Green Chair in Systems Biology Science to K.P. We gratefully acknowledge Dr. Joslyn Pribble and the Microbiology lab at Methodist Health System, without whom this work would not be possible.

