## Supplemental Text and Figures for "Vancomycin resistance in *Enterococcus faecium* from the Dallas, Texas area is conferred predominantly on pRUM-like plasmids"

**Running Head: Genomic analysis of Dallas VREfm isolates**

#### Supplementary Text

#### Supplemental Methods

**DNA isolation**. Genomic DNA was isolated as described previously (1). Briefly, cells were cultured in BHI broth at 37°C and pelleted from 1 mL overnight culture, washed with 1X TE buffer, and treated with 210 µL lysozyme solution at 37°C for 2 h prior to proteinase K treatment and column purification of gDNA using the Qiagen DNeasy Blood and Tissue kit per the manufacturer's instructions. The composition of lysozyme solution is 180 µL enzymatic lysis buffer (20 mM Tris Cl, 2 mM EDTA, 1.2% Triton X-100), 5 µL of 20 mg/mL lysozyme, 10 µL of 2.5 KU/mL mutanolysin and 15 µL of pre-boiled (95°C, ~5 min) 10 mg/mL RNase A.

Plasmids were isolated using the Qiagen Midiprep kit, adjusted according to the manufacturer's instructions to include low copy number plasmids. Briefly, VREfm cells were pelleted from 500 mL overnight culture and pretreated with 20 mL lysozyme solution prior to addition of buffer P1 and additional steps as directed by the manufacturer. Plasmid DNA was precipitated with 42 mL isopropanol and then column-purified.

**MinION sequencing**. VREfm isolates (n=47) were sequenced by Oxford Nanopore technology (2) as described (3) with minor adjustments. Briefly, for each sample, a total of 1.5 µg genomic DNA and midi-prepped plasmid were mixed in a ratio of 7:3 to prepare the library. The DNA mixture was mechanically sheared into 8 kb fragments with Covaris g-tube per manufacturer's instructions prior to library preparation with the ONT Ligation Sequencing Kit 1D (SQK-LSK108). Samples were multiplexed, and a total of 12 strains were sequenced using the Native Barcoding Kit (EXP-NBD103) in each flow cell. Libraries were base-called with MinKNOW (v3.5.5) to generate fastq and fast5 sequence reads.

**Illumina sequencing**. Library preparation and sequencing were performed by the Genome Center at the University of Texas at Dallas. Genomic DNA library was prepared using the Nextera Flex DNA library kit according to the manufacturer's instructions (Illumina). The resulting libraries were pooled and sequenced in the mid-output mode with paired-end reads of 2 x 150 bp on the NextSeq 500 sequencing platform.

**Hybrid genome assembly**. Processing of the reads and read assembly were completed on the operating system Ubuntu 18.04.4 LTS. The fastq sequences of the whole genome, generated by MinKNOW, were demultiplexed using the data processing toolkit Guppy (v 3.2.6) and then filtered to gather reads with q score >9 and length >1000 bp using Nanofilt (v2.5.0) (4). The adaptor sequences were trimmed from the filtered reads with Porechop (v 0.2.3) (https://github.com/rrwick/Porechop). The processed MinION reads were assembled with Illumina reads using Unicycler (v 0.4.7) (3) with the default setting "normal mode". Incomplete assemblies were manually completed as described in the "Unicycler tips for finishing genome" page (https://github.com/rrwick/Unicycler/wiki/Tips-for-finishing-genomes). Briefly, Bandage (v 0.8.1) (5) was used to visualize completion status of the assembly, and unassembled contig sequences were extracted. Using these unassembled contig sequences as baits, long reads from MinION sequences were gathered for incomplete regions using minimap2 (v 2.11-r797) (6) and in-Bandage BLAST search was performed with the long reads against the graph. If long reads supported the continuity of two unassembled contigs, then the Bandage graph editing function was used to duplicate, delete edge, and merge contigs. The complete assembly sequence was then saved from Bandage.

**Analysis of other closed circular DNA elements.** Eight closed circular DNA elements of varying sizes (3 to 72 kb) could not be classified by PlasmidFinder nor by the pMIX *rep* typing scheme described above. These 8 elements were observed in the genome assemblies of 8 different isolates (51-1, 52, 71-1, 111, 124-1, 124-2, 124-3, and 148). The completeness of the assemblies for these 8 isolates was visualized with Bandage (5). BLAST analysis was performed with the nucleotide sequences of these entities against the NCBI database to confirm the absence of contaminating sequence from other bacterial species. NCBI Conserved Domain Analysis (CDS) was used to identify plasmid (i.e. Rep) or pathogenicity island (PAI)-specific virulence genes (i.e. *esp, hyl* and *acm*) (7) as well as transposons or IS elements.

**Transcriptional activity of predicted toxin-antitoxin (TA) systems**. Transcriptional activity of the TA systems, $TA_{axe\text{-}txe}$ and $TA_{relE}$, was determined by RT-qPCR for the VREfm isolates 1 and 5. Total RNA was collected from overnight cultures as described previously (8). Briefly, cells were resuspended with 50 mg/mL lysozyme, 2.5 KU mutanolysin, and 500 µL IHB-1 (50 mM glucose, 25 mM Tris, 10 mM EDTA) and incubated at 37°C for 20 min. This pre-treated cell suspension was mixed with RNA-Bee (Tel-Test, Inc.) and lysed by bead beating in lysis matrix B tubes (MP Biomedicals). RNA was collected from the aqueous layer by ethanol precipitation. The collected RNA was treated with DNase (Roche), checked for rRNA integrity by agarose gel electrophoresis, and checked for DNA contamination by PCR. cDNA was synthesized from the total RNA with SuperScript Reverse Transcriptase III (Thermo Fisher) and random hexamers (Qiagen). qPCR was performed with AzuraQuantTM Green Fast qPCR Mix LoRox (Azura Genomics) according to the manufacturer’s instructions in the MyGo Pro real-time PCR instrument in biological triplicates. The expression level of the 4 genes of interest (*axe, txe, relE*-toxin, and *relE*-antitoxin) were measured relative to the expression level of the housekeeping gene 16S rRNA. The primer sequences used for qPCR are provided in **Dataset S1C.**

#### Supplemental Results

**Analysis of unidentified circular entities in the genome assemblies.** Eight closed circular entities of various sizes (2.6 to 72 kb) observed in the genome assemblies of 8 different isolates (51-1, 52, 71-1, 111, 124-1, 124-2, 124-3, and 148) could not be classified by PlasmidFinder or the pMIX typing scheme (**Dataset S1H**). All had at least partial sequence identity to other *E. faecium* chromosomes or megaplasmids present in the NCBI public sequence database, suggesting that these sequences are not the result of contamination.

Complete genome assemblies were achieved for 4 out of the 8 isolates (111, 124-1, 124-3, and 148. The isolates 71-1 and 51-1 had incomplete chromosome assemblies. The isolates 52 and 124-2 had incomplete megaplasmid assemblies (**Dataset S1H**). It is therefore possible that these unclassified DNA entities arose from assembly errors in isolates 71-1, 51-1, 52, and 124-2, although their circular instead of linear assemblies argue against this.

The 8 elements did not encode conserved domains specific for plasmid replication or virulence genes of the *E. faecium* PAI (**Dataset S1I**). However, each harbored 2 to 6 transposable elements (TE), and none of them were identical to (i.e. repetitive with) sequences elsewhere in the same genome. Therefore we conclude that these 8 circular DNA elements are likely to be excision products of genetic islands and transposable elements.

**Analysis of plasmid replicon distribution in commonly observed STs.** To evaluate whether plasmid replicons were enriched among specific lineages, the percentage presence/absence of specific *rep* genes was quantified among the isolates belonging to 4 STs that included at least 5 isolates (ST17, ST18, ST1703 and ST664). The other 6 STs (ST262, ST1383, ST612, ST412, ST80, and ST736) were not included in this analysis because they were represented by <5 isolates (**Dataset S1G**). Chromosomally integrated *rep* genes *repUS12* and *repUS43,* encoding the protein motifs Rep_1 and Rep_trans, were observed in 45% and 22% of the isolates, respectively, and were absent from isolates belonging to ST17, ST18 and ST1383 (**Dataset S1G**). The *rep14* family was present in 80% and 90% of ST17 and ST18 isolates, respectively, but was absent in ST1703 and ST664 isolates. The *rep18* family plasmid was present in ST17 (80%) and ST664 (100%) isolates but absent in ST1703 (0%) and rare in ST18 (30%). In summary, the *rep* genes *repUS12, repUS43*, *rep18* and *rep14* had an ST-specific distribution among the isolates. No conclusion could be drawn regarding other *rep* genes and STs due to low representation of some STs in the strain collection.

**Detailed analysis of resistance genes detected.**

**Aminoglycoside resistance**

***i) aac(6')-aph(2"):*** The bifunctional enzyme 6'-aminoglycoside acetyltransferase/2"-aminoglycoside phosphotransferase*,* sharing 99.4 to 100% nucleotide sequence identity with the reference gene (accession number M13771) was detected in 83.3% (39 out of 47) of the Dallas VREs (**Dataset S1L**). Mostly plasmid-borne, this gene was encoded in the megaplasmid (*repUS15*) in 92% (36 out of 39) of the cases. The VRE isolate 66 (ST18) was the only one to carry this gene in the pRUM-like resistance plasmid. Chromosomally-encoded *aac(6')-aph(2")* was observed in the isolate 163-1. The position of *aac(6')-aph(2")* was uncertain in the isolate 148 since the genome was not completely closed. This gene was absent in 16.6% (8 out of 47) of the isolates belonging to ST18 (isolate 5 and 144-1), ST612 (isolate 51-4) and ST1703 (isolate 111, 121, 137, 154-1 and 158).

***ii) ant(6)-Ia:***

The gene *ant(6)-Ia* (Accession number AF330699) is the aminoglycoside nucleotidyltransferase that represents a cluster of 3 resistance genes *aadE* (6'-aminoglycoside adenylyltransferase), *sat4* (streptothricin acetyltransferase) and *aphA* (3' aminoglycoside phosphotransferase) conferring resistance against streptomycin, streptothricin and kanamycin (9). Sharing 99.2 to 100% nucleotide sequence identity with the reference gene AF330699, the *ant(6)-Ia* gene cluster was observed in 80.8% (38 out of 47) of the VRE isolates. The pRUM-like plasmid harbored this gene in 92% (35 out of 38) of the isolates while in the remaining 3 isolates it was chromosomally encoded, among which 53-1 and 53-2 encoded the resistance gene on the chromosomally integrated *rep18* plasmid, while the isolate 91 lacked *rep18* in its genome.

This gene was absent in 19.1% (9 out of 47) of the isolates, which were of different genetic backgrounds: ST80 (isolates 16-2 and 113), ST17 (isolate 93-3), ST18 (isolates 97-1, 124-1, 124-2 and 124-3), and ST1703 (isolate 111 and 154-1). The plasmid *rep* group *rep18* was absent in 7 of these isolates. An indication of a correlation between *rep18* plasmids and *ant(6)-Ia* can be drawn from this observation.

**Macrolide resistance genes were encoded in both plasmids and the chromosome**

Resistance against macrolide antibiotics including erythromycin, lincosamide, azithromycin, tylosin and quinupristin were conferred by a combination of mostly *erm(B)* and *msr(C)* genes among the Dallas VREs. The presence of the resistance genes *erm(T)* and *lnu* were rarely observed among these isolates. The gene *msr(C),* sharing 99% nucleotide sequence identity with the reference gene AY004350, was present in all 47 isolates. It was encoded in the chromosome of 45 out of the 47 isolates; in the remaining 2 isolates the position of this gene could not be conclusively stated due to incomplete assembly. The gene *erm(B),* sharing 99.9% to 100% nucleotide sequence identity with the reference gene AF299292, was detected in 85.4% (41 out of 47) of the Dallas VREs. The pRUM-like plasmid encoded *erm(B)* in 92.7% (38 out of 41) of the isolates while in 7.3% (3 out of 41) of the isolates the *erm(B)* gene was encoded in the chromosome, despite the fact that the pRUM-like plasmid was present in these strains. This gene was absent in 12.7% (6 out of 47) VRE isolates of different genetic backgrounds: ST80 (isolate

16-2 and 113), ST1703 (isolate 111, 154-1), ST18 (isolate 162), and ST262 (isolate 163-1). The rare chromosomally-encoded gene *erm(T)* was observed in only 3 out of 47 of the isolates among which 2 isolates (16-2 and 163-1) lacked *erm(B)*. The isolate 16-1 was the only one harboring both *erm(B)* and *erm(T)*. These data suggest that *ermB* is the predominant gene conferring erythromycin resistance in the isolates and mainly resides on the pRUM-like resistance plasmid.

**Tetracycline resistance was common among the Dallas VREfm isolates**

The chromosomally encoded tetracycline resistance genes *tet(M)* and *tet(L)* identified among the Dallas VREfm shared 97.5% and 99.9% sequence identity, respectively, with the reference genes (accession no EU182585 and M29725, respectively). All of the isolates, except 100-1 and 100-2, harbored *tet(M)*, and *tet(L)* was also present in 21 of 47 of isolates. The isolates 100-1 and 100-2 lack both *tet(M)* and *tet(L)* and theoretically should be sensitive to the drug. Interestingly, isolates belonging to ST17 or ST18 did not code for *tet(L).* It can be concluded that *tet(M)* is the major tetracycline resistance gene among the Dallas VREfm isolates.

**Trimethoprim resistance was rare among the Dallas VREfm isolates.**

The trimethoprim resistance gene *dfrG*, identified in 13 out of 47 of the Dallas VREs, shared 100% nucleotide sequence identity with the reference gene (accession No AB205645). This resistance gene was encoded in the pMG1 plasmid in only 3 out of 13 of the isolates belonging to ST18 (isolates 66 and 144-1) and ST612 (isolate 51-4). These three isolates were the only ones among the Dallas VREfm collection that harbored the pMG1 plasmid at all. Chromosomally encoded *dfrG* was observed in the rest of the *dfrG*-encoding isolates (10 out of 13), which belonged to a range of genetic backgrounds: ST1703 (isolates 111, 121 and 137), ST736 (isolates 9-1 and 9-2), ST412 (isolate 17-1) and ST262 (isolate 163-1). None of the isolates belonging to ST17 harbored this resistance gene.

**Specific structural variations of the Dallas Tn*1546* elements:**

**The 3' but not 5' boundary of the Tn*1546* was conserved among the Dallas isolates.** Each of the 17 Tn*1546* groups preserved the $IR_R$ boundary of Tn*1546* but lacked the $IR_L$ boundary (**Figure 7**). The only exception was the isolate 17-1 (group BBCD) that lost both of the IR boundaries due to insertion of IS*1216*. The complete or partial deletion of ORF1 (transposase) and ORF2 (resolvase) was observed in all of the Tn*1546* groups. This might decrease the likelihood of these Tn*1546* to disseminate to other isolates without a plasmid as a vehicle.

**IS*1251* and IS*1216* were the most common IS elements.** IS*1251* was present in 84.4% (38 of 45) of the isolates (**Table 3**). It was inserted at Tn*1546* position 5820 in the *vanS*-*vanH* intergenic region, in reverse orientation to the *van* genes (**Figure 7**). The IS*1251* generated an 8 bp direct repeat (5'-AAAATTAT-3') in the regions where it was inserted. IS*1216* was inserted at either the 5' end of Tn*1546* in 71.1% (32 of 45) or in the intergenic region of *vanX*-*vanY* in 17.8% (8 of 45) of the isolates.

**The *van* genes and *vanR* and *vanH* promoters were intact in most isolates, despite IS element insertions.** IS*1251* insertion at Tn*1546* position 5820 did not disrupt the two 12 bp VanR binding sites located at positions 5846 and 5879 (**Figure S2**). The 70 bp VanR-interacting region (-110 to -41) from positions 5935 to 6004 is also intact. The 197 bp long proposed *vanR* promoter region (-173 to +24) is located at the position 3,803 to 3,999 bp of the Tn*1546*. The isolate 137 (Tn*1546* group BCK) has an IS*Efa16* inserted at the position 3664 to 3763 that was responsible for a 99 bp deletion of resolvase at 3' region but the *vanR* promoter region remained uninterrupted. The *vanR* promoter region including the 12 bp consensus VanR binding site is disrupted in the isolates 51-1, 87-1, 93-1, 93-2, 155, 160 from group J because of the insertion of IS*1216* at position 3914. The *vanR* promoter region for other isolates remained uninterrupted.

Deletion of *vanR* and *vanS* was observed in the isolate 91 (variant group D1). Expression of the *vanHAX* genes may be constitutive instead of inducible in this isolate. No mutations were identified in *vanR* and *vanS* in the rest of the groups.

**Two point mutations in *vanHAX* were fixed in the population.** All isolates harbored a point mutation (T7658C) in *vanA* that generated a valine to alanine substitution (**Table 3**) and a second point mutation (G8234T) in *vanX* that generated a lysine to asparagine substitution. 62.2% (28 of 45) of the isolates harbored a third point mutation (C9692T) in *vanY* that was also present in the reference strain ATCC® 700221™. This point mutation generates a proline to leucine substitution in VanY. Isolates 53-1 and 53-2 (group BC10) had point mutations C6513A and G6514A in *vanH* that generated an arginine to lysine substitution. Isolate 144-1 harbored a unique G9566A point mutation in *vanY* that conferred a glycine to glutamic acid substitution. No mutations in *vanZ* were observed in any isolates.

**Summary.** A total of 17 variations of the Tn*1546* region, defined by the presence of different IS elements, point mutations, and deletions, were observed among the Dallas isolates. In absence of a strict guideline for the nomenclature of the Tn*1546* variants, a recent one described in 2017 (10) was followed in this study. A slight variation in Tn*1546* nomenclature was utilized in 2016 where variants with IS*1251* were designated as “type F” or “US hospital type” (11), which are designated as group “C” in the current study. Three newly identified IS elements (IS*Efa16,* IS*Efa17,* and IS*Efa18*) within the Tn*1546* were observed in this study. Moreover, two point mutations within Tn*1546*, T7658C and G8234T, which were ubiquitous among the Dallas VREfm isolates, were reported previously but never in the same Tn*1546* group (10). The point mutation T7658C was reported in the Tn*1546* groups BC5, BC1, BC2, BC3, and BC4 while G8234T was reported in groups A2, C1, C2, BB1, BB2, BC1, BC2, BC3, BC4, BC5, BBBI1, and BBBI2 (10).

### DNA alignment

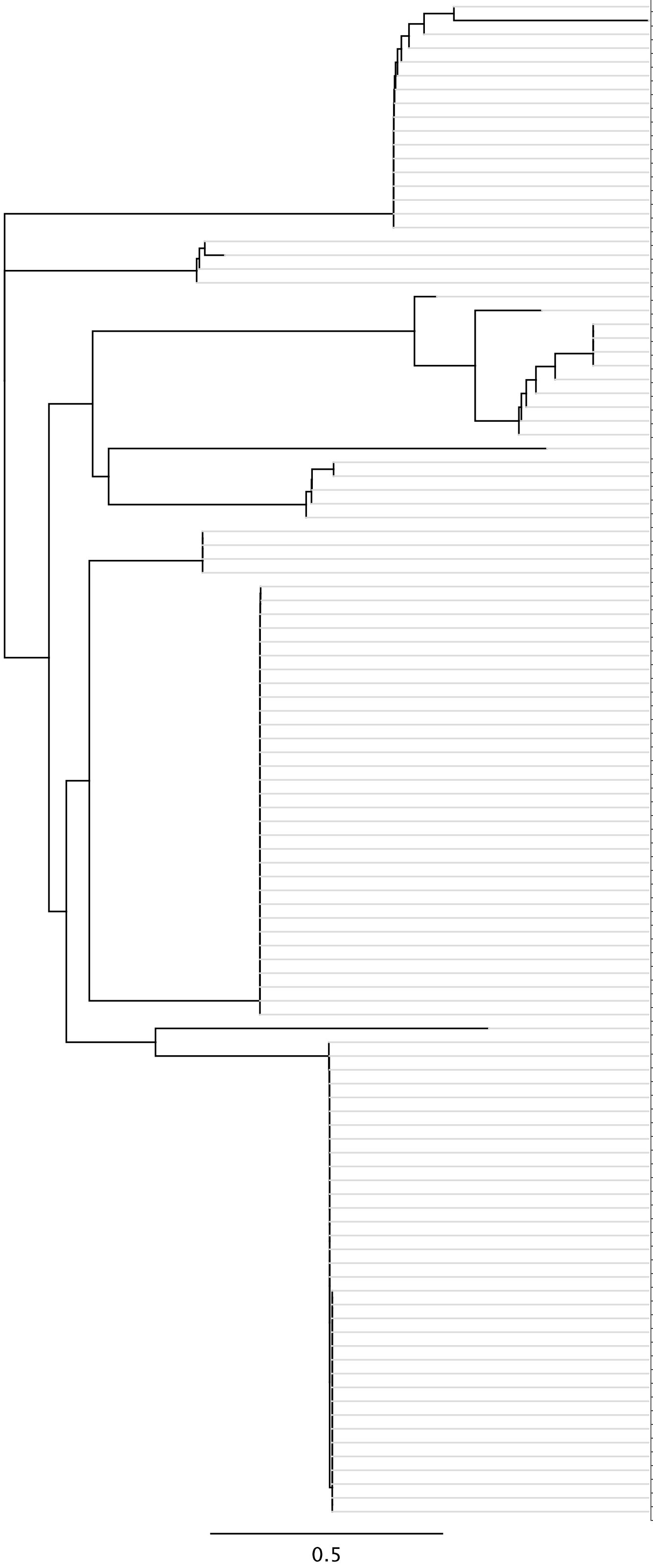


| Isolate | Plasmid Size (kb) | *rep* gene | Plasmid Group |
|---|---|---|---|
| 9-1 | 4.3 | rep | pMI3 |
| 97-1 | 3.6 | | |
| 9-2 | 4.3 | | |
| 43-1 | 4.3 | | |
| 52 | 4.3 | | |
| 66 | 4.3 | | |
| 71-1 | 4.3 | | |
| 71-2 | 4.3 | | |
| 97-1 | 4.3 | | |
| 107-1 | 4.3 | | |
| 124-1 | 4.3 | | |
| 124-2 | 4.3 | | |
| 124-3 | 4.3 | | |
| 131-2 | 4.3 | | |
| 137 | 4.3 | | |
| 160 | 4.3 | | |
| 163-1 | 4.3 | | |
| 9-1 | 3 | repB | pMI7 |
| 17-1 | 3 | | |
| 9-2 | 3 | | |
| 137 | 3 | | |
| 51-4 | 4.3 | rep | pMI4 |
| 100-1 | 11 | | |
| 17-1 | 4.3 | | |
| 52 | 4.3 | | |
| 93-2 | 4.3 | | |
| 111 | 4.3 | | |
| 51-1 | 4.3 | | |
| 55 | 4.3 | | |
| 100-2 | 8.4 | | |
| 155 | 4.3 | | |
| 160 | 4.3 | | |
| 5 | 70 | Rol_rep_N | pMI9 |
| 16-1 | 104 | repB | pMI5 |
| 16-2 | 90 | | |
| 51-1 | 83 | | |
| 51-4 | 83 | | |
| 144-1 | 87 | | |
| 16-1 | 4.4 | repB | pMI6 |
| 16-2 | 4.4 | | |
| 51-4 | 4.4 | | |
| 113 | 4.4 | | |
| 1 | 6.3 | repB | pMI2 |
| 93-3 | 6.3 | | |
| 9-1 | 6.3 | | |
| 9-2 | 6.3 | | |
| 16-1 | 6.3 | | |
| 16-2 | 6.3 | | |
| 43-1 | 6.3 | | |
| 51-1 | 6.3 | | |
| 53-1 | 6.3 | | |
| 53-2 | 6.3 | | |
| 57 | 6.3 | | |
| 66 | 8 | | |
| 71-1 | 6.3 | | |
| 71-2 | 6.3 | | |
| 91 | 6.3 | | |
| 93-2 | 6.3 | | |
| 103 | 6.3 | | |
| 107-1 | 6.3 | | |
| 111 | 6.3 | | |
| 124-1 | 6.3 | | |
| 124-2 | 6.3 | | |
| 124-3 | 6.3 | | |
| 131-2 | 6.3 | | |
| 137 | 6.3 | | |
| 144-1 | 6.3 | | |
| 148 | 6.3 | | |
| 154-1 | 6.3 | | |
| 155 | 6.3 | | |
| 158 | 6.3 | | |
| 160 | 6.3 | | |
| 161 | 6.3 | | |
| 163-1 | 6.3 | | |
| 51-4 | 2 | repB | pMI8 |
| 5 | 56 | repA | pMI1 |
| 9-1 | 54 | | |
| 9-2 | 54 | | |
| 51-1 | 62 | | |
| 52 | 54 | | |
| 55 | 54 | | |
| 93-2 | 62 | | |
| 97-1 | 56 | | |
| 17-1 | 56 | | |
| 100-1 | 56 | | |
| 100-2 | 56 | | |
| 121 | 59 | | |
| 137 | 61 | | |
| 154-1 | 23 | | |
| 155 | 62 | | |
| 158 | 23 | | |
| 160 | 62 | | |
| 163-1 | 23 | | |
| 1 | 55 | | |
| 16-1 | 60 | | |
| 43-1 | 55 | | |
| 51-4 | 60 | | |
| 53-1 | 55 | | |
| 53-2 | 55 | | |
| 57 | 57 | | |
| 66 | 57 | | |
| 71-2 | 57 | | |
| 71-1 | 57 | | |
| 91 | 55 | | |
| 93-3 | 57 | | |
| 124-1 | 59 | | |
| 124-2 | 59 | | |
| 124-3 | 59 | | |
| 131-2 | 55 | | |
| 144-1 | 56 | | |

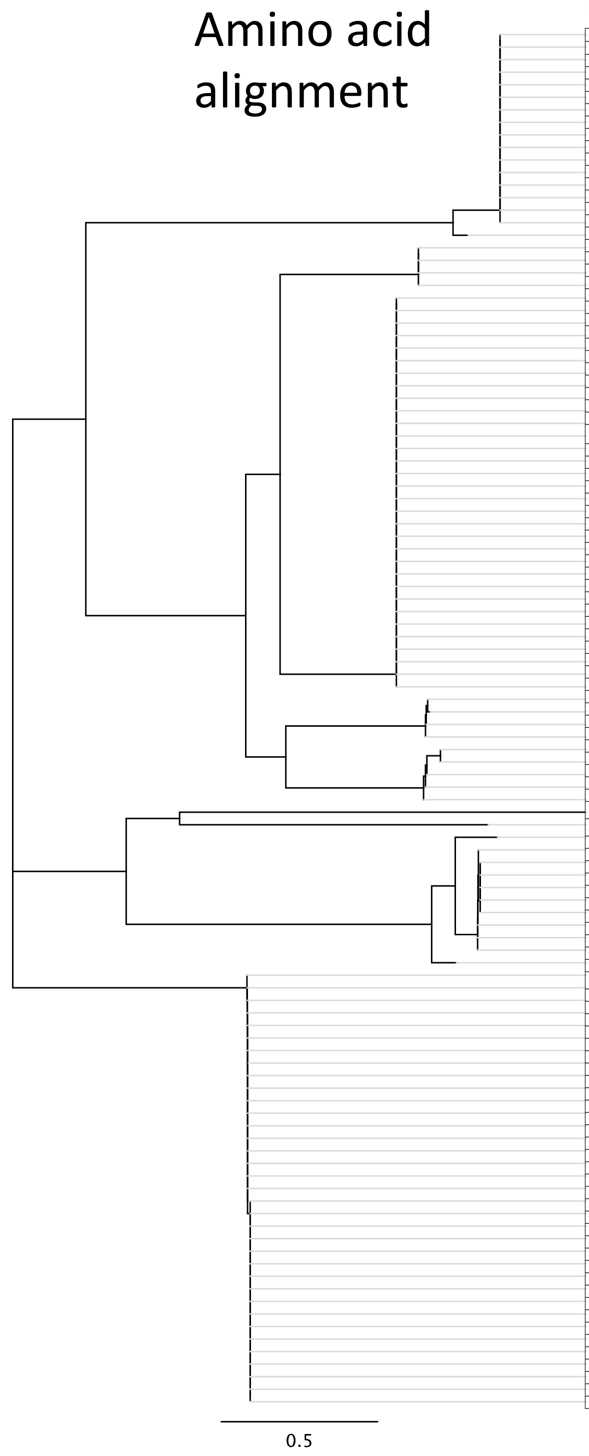


| Isolate | Plasmid Size (kb) | *rep* gene | Plasmid Group |
|---|---|---|---|
| 9-1 | 4.3 | rep | pMI3 |
| 9-2 | 4.3 | | |
| 43-1 | 4.3 | | |
| 52 | 4.3 | | |
| 66 | 4.3 | | |
| 71-1 | 4.3 | | |
| 71-2 | 4.3 | | |
| 97-1 | 4.3 | | |
| 107-1 | 4.3 | | |
| 124-1 | 4.3 | | |
| 124-2 | 4.3 | | |
| 124-3 | 4.3 | | |
| 131-2 | 4.3 | | |
| 137 | 4.3 | | |
| 160 | 4.3 | | |
| 163-1 | 4.3 | | |
| 97-1 | 3.6 | | |
| 16-1 | 4.4 | repB | pMI6 |
| 16-2 | 4.4 | | |
| 51-4 | 4.4 | | |
| 113 | 4.4 | | |
| 1 | 6.3 | repB | pMI2 |
| 93-3 | 6.3 | | |
| 9-1 | 6.3 | | |
| 9-2 | 6.3 | | |
| 16-1 | 6.3 | | |
| 16-2 | 6.3 | | |
| 43-1 | 6.3 | | |
| 51-1 | 6.3 | | |
| 53-1 | 6.3 | | |
| 53-2 | 6.3 | | |
| 57 | 6.3 | | |
| 66 | 8 | | |
| 71-1 | 6.3 | | |
| 71-2 | 6.3 | | |
| 91 | 6.3 | | |
| 93-2 | 6.3 | | |
| 103 | 6.3 | | |
| 107-1 | 6.3 | | |
| 111 | 6.3 | | |
| 124-1 | 6.3 | | |
| 124-2 | 6.3 | | |
| 124-3 | 6.3 | | |
| 131-2 | 6.3 | | |
| 137 | 6.3 | | |
| 144-1 | 6.3 | | |
| 148 | 6.3 | | |
| 154-1 | 6.3 | | |
| 155 | 6.3 | | |
| 158 | 6.3 | | |
| 160 | 6.3 | | |
| 161 | 6.3 | | |
| 163-1 | 6.3 | | |
| 9-1 | 3 | repB | pMI7 |
| 17-1 | 3 | | |
| 9-2 | 3 | | |
| 137 | 3 | | |
| 16-1 | 104 | repB | pMI5 |
| 16-2 | 90 | | |
| 51-1 | 83 | | |
| 51-4 | 83 | | |
| 144-1 | 87 | | |
| 51-4 | 2 | repB | pMI8 |
| 5 | 70 | Rol_rep_N | pMI9 |
| 51-4 | 4.3 | rep | pMI4 |
| 17-1 | 4.3 | | |
| 51-1 | 4.3 | | |
| 55 | 4.3 | | |
| 100-2 | 8.4 | | |
| 155 | 4.3 | | |
| 160 | 4.3 | | |
| 52 | 4.3 | | |
| 93-2 | 4.3 | | |
| 111 | 4.3 | | |
| 100-1 | 11 | | |
| 5 | 56 | repA | pMI1 |
| 9-1 | 54 | | |
| 9-2 | 54 | | |
| 51-1 | 62 | | |
| 52 | 54 | | |
| 55 | 54 | | |
| 93-2 | 62 | | |
| 17-1 | 56 | | |
| 97-1 | 56 | | |
| 100-1 | 56 | | |
| 100-2 | 56 | | |
| 121 | 59 | | |
| 137 | 61 | | |
| 154-1 | 23 | | |
| 155 | 62 | | |
| 158 | 23 | | |
| 160 | 62 | | |
| 163-1 | 23 | | |
| 1 | 55 | | |
| 16-1 | 60 | | |
| 43-1 | 55 | | |
| 51-4 | 60 | | |
| 53-1 | 55 | | |
| 53-2 | 55 | | |
| 57 | 57 | | |
| 66 | 57 | | |
| 71-2 | 57 | | |
| 71-1 | 57 | | |
| 91 | 57 | | |
| 93-3 | 55 | | |
| 124-1 | 5759 | | |
| 124-2 | 59 | | |
| 124-3 | 59 | | |
| 131-2 | 55 | | |
| 144-1 | 56 | | |

**Figure S1. Alignment of replicon sequences in 110 pMIX plasmids**. Nucleotide (*rep*) and amino acid sequences (Rep) were aligned with MUSCLE, and neighbor joining trees are shown. A total of 9 plasmid groups (pMI1 to pMI9) were assigned based on the clustering of sequences within the trees.

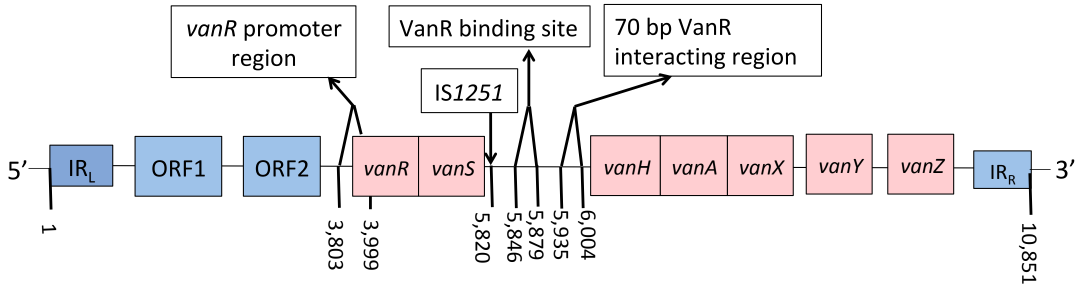


**Figure S2. Tn*1546* structure and regulatory regions.** The Tn*1546* structure consisting of IR$_L$ (inverted repeat, left), ORF1 (transposase), ORF2 (resolvase), regulatory and resistance genes (*vanR, vanS, vanH, vanA, vanX, vanY,* and *vanZ*), and IR$_R$ (inverted repeat, right) are delineated. The 5' region of IR$_L$ marks the starting position 1 while the 3' region of the IR$_R$ boundary marks the end position 10,851 of the Tn*1546.* The positions of the *vanR* promoter region at 3,803 and 3,999 (-173 and +24 regions), the VanR binding site at positions 5,846 and 5,879, and the 70 bp VanR interacting region at positions 5,935 and 6,004 are indicated. The figure is not drawn to scale.

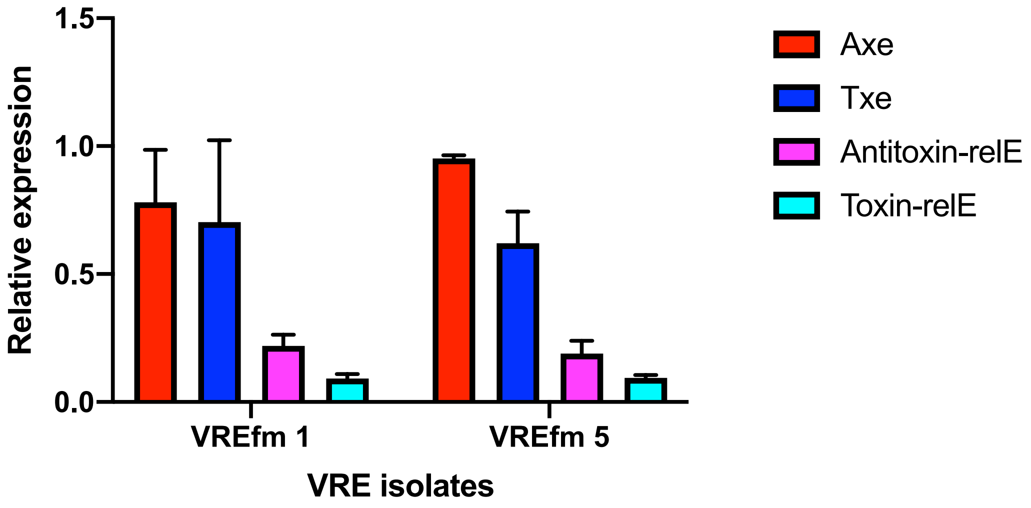


**Figure S3. Toxin-antitoxin (TA) systems are expressed during culture in laboratory medium**. mRNA expression levels of antitoxin and toxin components of $TA_{axe\text{-}txe}$ and $TA_{relE}$ were quantified by RT-qPCR in the isolates VREfm1 and VREfm5. The y-axis shows the expression levels relative to the housekeeping gene 16S rRNA.

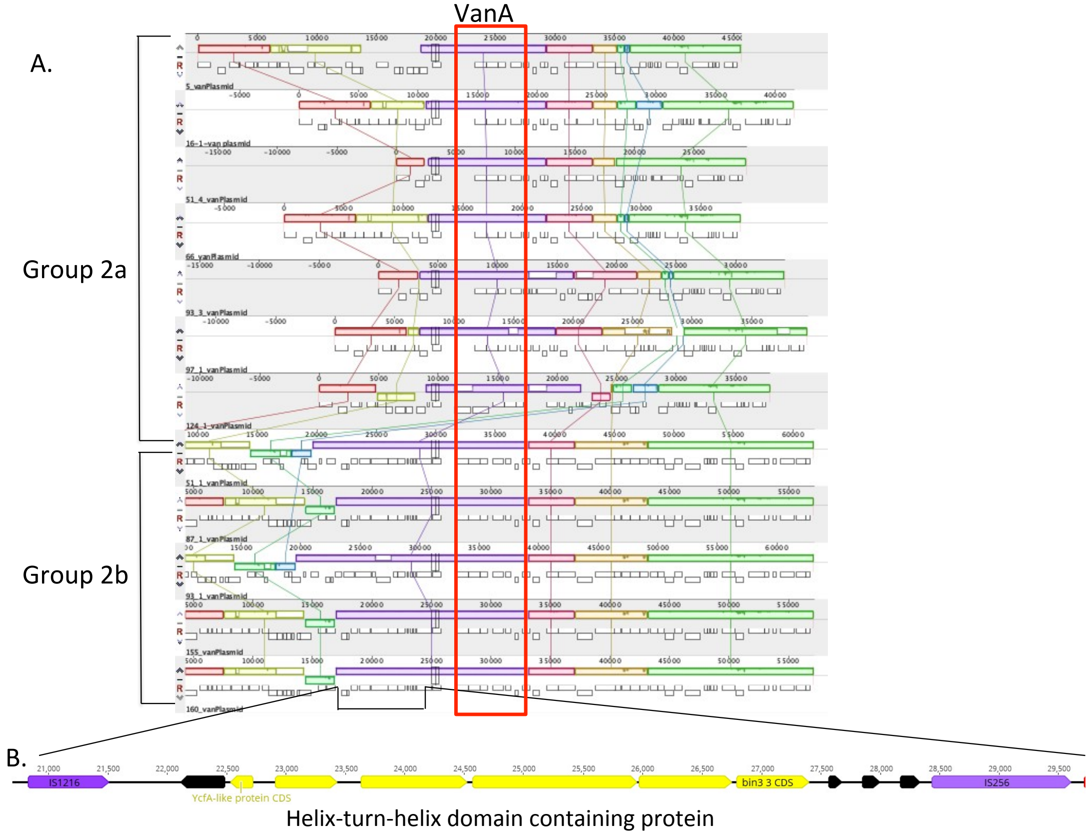


**Figure S4. Unique region in Group 2b pRUM-like plasmids.** A) Alignment of the pRUM-like plasmids from group 2a and 2b. The alignment was generated by progressive Mauve algorithm from Geneious. The *van* gene cluster is highlighted with a red box. The region containing the helix-turn-helix XRE family-like DNA binding protein is indicated at the base of the alignment. This region is unique to the group 2b resistance plasmids. **B**. Sketch of the region unique to group 2b is presented. Arrows indicate coding regions. Yellow arrows indicate predicted helix-turn-helix domain-containing proteins. Black arrows indicate hypothetical genes. IS elements are represented by purple arrows.

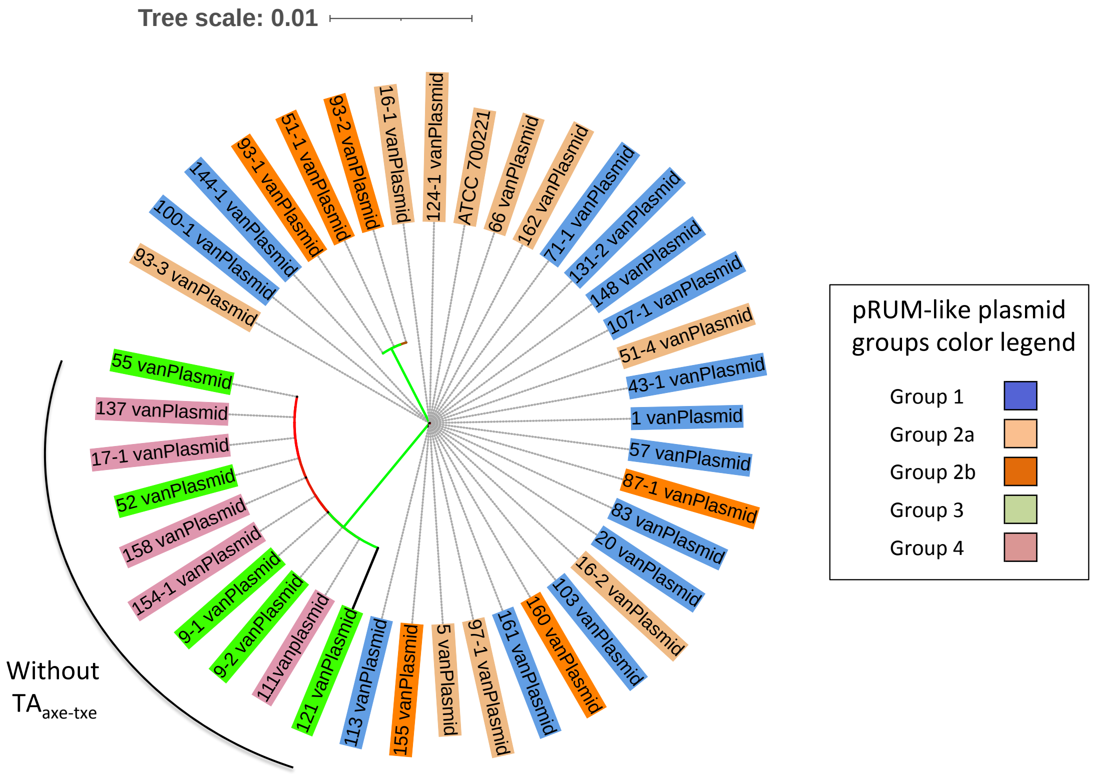


**Figure S5. pRUM-like plasmid core genome phylogeny.** The RAxML phylogeny is based on an alignment of SNPs in the 7 core genes found in 40 pRUM-like plasmids. The tips of the tree are labeled with *E. faecium* isolate names. The tree tip labels are color-coded according to the plasmid groups 1, 2a, 2b, 3 and 4. The branches colored green and red are supported by a bootstrap value of >70, and <30 respectively.

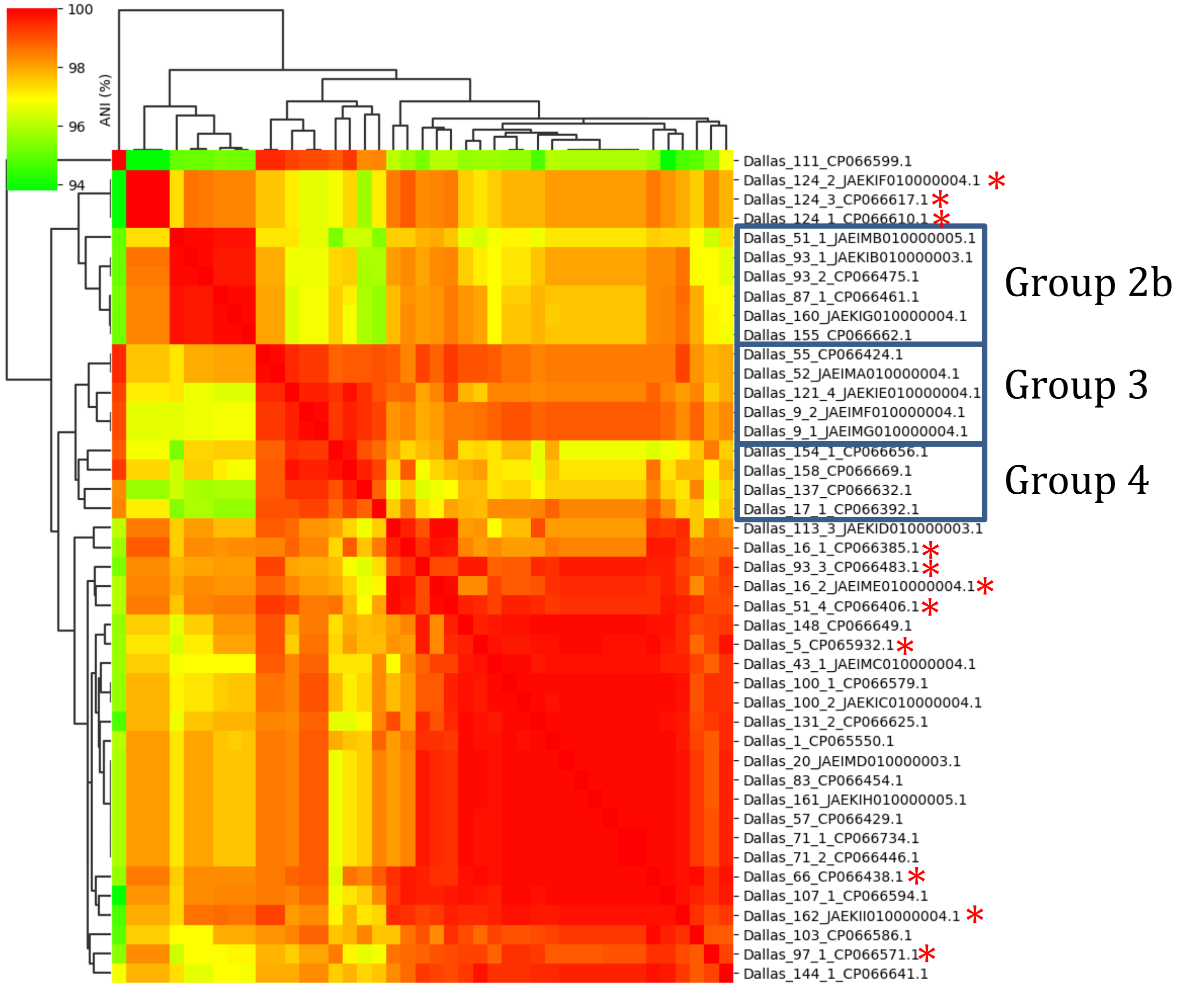


**Figure S6. ANI analysis of pRUM-like plasmids.** Percent ANI is color-coded as shown in the legend. Plasmid groups 2b, 3, and 4 (with the exception of the pRUM-like plasmid from isolate 111, at top of figure) are shown in boxes. Plasmids belonging to group 2a are denoted with red stars. All other plasmids belong to group 1.

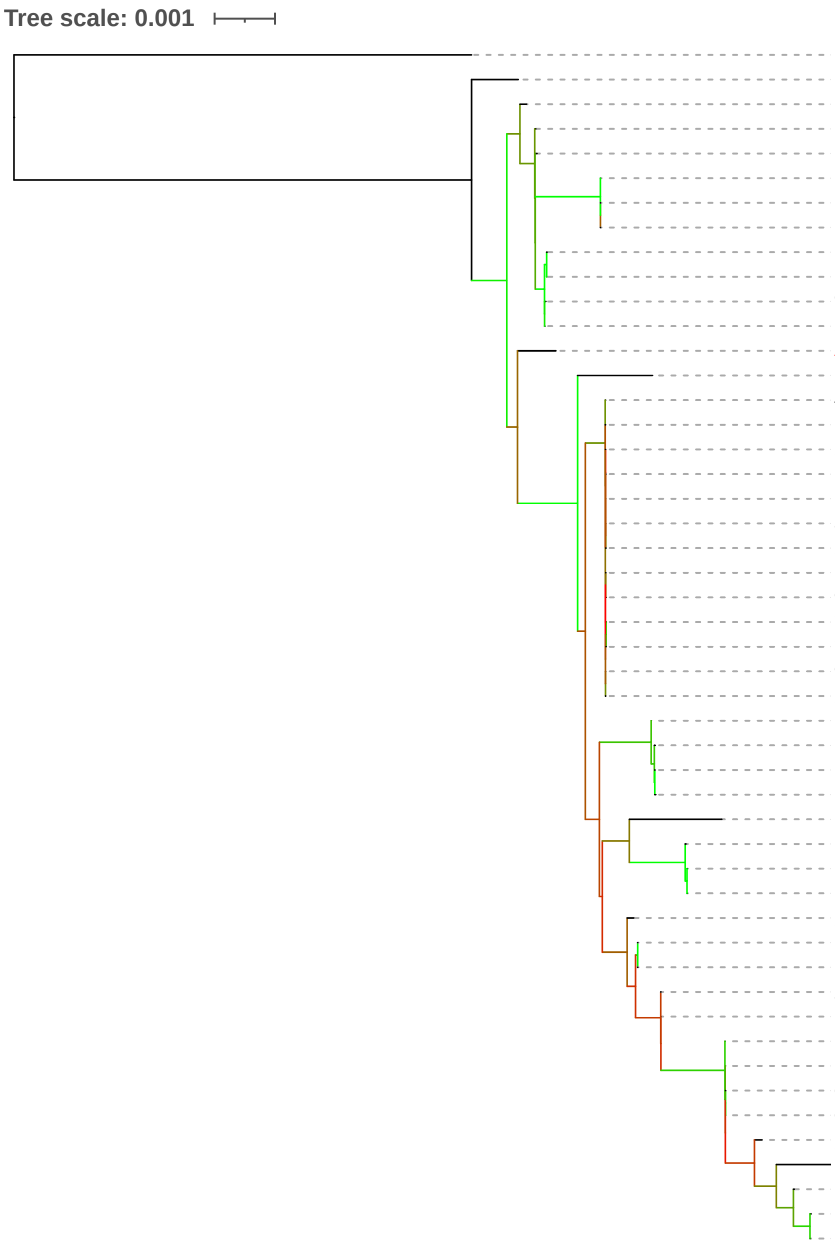


| Strain | MLST | Van-plasmid type 1 | 2a | 2b | 3 | 4 | Tn*1546* type C2 | C3 | D1 | BC(2-11) | BBC/BCB | BCCL/BCL | BCM | BCK | J | BBCD |
|---|---|---|---|---|---|---|---|---|---|---|---|---|---|---|---|---|
| 1,231,501 | | N/A | | | | | N/A | | | | | | | | | |
| 163-1 | | N/A | | | | | N/A | | | | | | | | | |
| 162 | | | ❖ | | | | | | | + | | | | | | |
| 66 | | | ❖ | | | | | | | + | | | | | | |
| 144-1 | | ❖ | | | | | | | | + | | | | | | |
| 124-3 | | | ❖ | | | | | | | | | + | | | | |
| 124-2 | | | ❖ | | | | | | | | | + | | | | |
| 124-1 | | | ❖ | | | | | | | | | + | | | | |
| 100-2 | | ❖ | | | | | | | | + | | | | | | |
| 100-1 | | ❖ | | | | | | | | + | | | | | | |
| 97-1 | | | ❖ | | | | | | | | + | | | | | |
| 5 | | | ❖ | | | | | | | + | | | | | | |
| ATCC 700221 | | | ❖ | | | | + | | | | | | | | | |
| 51-4 | | | ❖ | | | | | | | + | | | | | | |
| 43-1 | | ❖ | | | | | | | | | + | | | | | |
| 131-2 | | ❖ | | | | | | | | + | | | | | | |
| 53-2 | | N/A | | | | | | | | ✓ | | | | | | |
| 148 | | ❖ | | | | | | | | + | | | | | | |
| 1 | | ❖ | | | | | | | | + | | | | | | |
| 83 | | ❖ | | | | | | | | + | | | | | | |
| 53-1 | | N/A | | | | | | | | ✓ | | | | | | |
| 103 | | ❖ | | | | | | | | + | | | | | | |
| 91 | | N/A | | | | | | | ✓ | | | | | | | |
| 71-2 | | ❖ | | | | | | | | + | | | | | | |
| 71-1 | | ❖ | | | | | | | | + | | | | | | |
| 93-3 | | | ❖ | | | | | | | | | | + | | | |
| 57 | | ❖ | | | | | | | | + | | | | | | |
| 137 | | | | | | ❖ | | | | | | | | + | | |
| 111 | | | | | | ❖ | Fragmented Tn1546 | | | | | | | | | |
| 154-1 | | | | | | ❖ | | | | | + | | | | | |
| 158 | | | | | | ❖ | | | | | + | | | | | |
| 17-1 | | | | | | ❖ | | | | | | | | | | + |
| 113 | | ❖ | | | | | | | | | + | | | | | |
| 16-1 | | | ❖ | | | | | | | | + | | | | | |
| 16-2 | | | ❖ | | | | | | | | + | | | | | |
| 52 | | | | | ❖ | | | + | | | | | | | | |
| 9-1 | | | | | ❖ | | | | | + | | | | | | |
| 9-2 | | | | | ❖ | | | | | + | | | | | | |
| 87-1 | | | | ❖ | | | | | | | | | | | + | |
| 155 | | | | ❖ | | | | | | | | | | | + | |
| 160 | | | | ❖ | | | | | | | | | | | + | |
| 51-1 | | | | ❖ | | | | | | | | | | | + | |
| 93-2 | | | | ❖ | | | | | | | | | | | + | |
| 93-1 | | | | ❖ | | | | | | | | | | | + | |
| 55 | | | | | ❖ | | | + | | | | | | | | |
| 121 | | | | | ❖ | | | | | + | | | | | | |
| 20 | | ❖ | | | | | | | | + | | | | | | |
| 107-1 | | ❖ | | | | | | | | + | | | | | | |
| 161 | | ❖ | | | | | | | | + | | | | | | |

**Figure S7. Core genome phylogeny, MLST, pRUM-like plasmid groups, and Tn*1546* types.** The tree is as shown in Figure 1. Presence of pRUM-like plasmids of 5 different types (1, 2a, 2b, 3 and 4) are indicated by the symbol (❖). The isolates 163-1 and the reference *E. faecium* 1,231,501 lack *vanA*-encoding Tn*1546*. Isolates 53-1, 53-2, and 91 have chromosomally integrated Tn*1546.* In all of these cases a variant of pRUM-like plasmid exists in the genome but they are not denoted as van-plasmids, except for *E. faecium* 1,231,501 for which plasmid information is unavailable due to its incomplete genome assembly. Structural variants of Tn*1546* are indicated with the symbol (✚). The symbol (✔) indicates chromosomally integrated Tn*1546.* The absence of Tn*1546* in the isolates 163-1 and 1,231,501 is indicated by "N/A". The *vanA* operon was encoded in two different contigs in isolate 111 therefore the Tn*1546* was considered to be fragmented.
